# MAVISp: A Modular Structure-Based Framework for Protein Variant Effects

**DOI:** 10.1101/2022.10.22.513328

**Authors:** Matteo Arnaudi, Mattia Utichi, Kristine Degn, Matteo Tiberti, Ludovica Beltrame, Karolina Krzesińska, Pablo Sánchez-Izquierdo Besora, Eleni Kiachaki, Simone Scrima, Laura Bauer, Katrine Meldgård, Anna Melidi, Lorenzo Favaro, Anu Oswal, Guglielmo Tedeschi, Terézia Dorčaková, Alberte Heering Estad, Joachim Breitenstein, Jordan Safer, Paraskevi Saridaki, Valentina Sora, Francesca Maselli, Philipp Becker, Jérémy Vinhas, Alberto Pettenella, Matteo Lambrughi, Claudia Cava, Anna Rohlin, Mef Nilbert, Sumaiya Iqbal, Peter Wad Sackett, Burcu Aykac Fas, Elena Papaleo

**Affiliations:** Cancer Structural Biology, Danish Cancer Institute, 2100, Copenhagen, Denmark; Cancer Systems Biology, Section for Bioinformatics, Department of Health and Technology, Technical University of Denmark, 2800, Lyngby, Denmark; Institute of Molecular Bioimaging and Physiology, National Research Council (IBFM-CNR), Via F. Cervi 93, Segrate-Milan, 20054, Milan, Italy; Bioinformatic and Computational Biology group, the Center for Development of Therapeutics, Broad Institute of MIT and Harvard, 415 Main St, Cambridge, MA 02142; Laboratoire de Biochimie Théorique, CNRS (UPR9080), Université Paris Cité, F-75005 Paris, France; Department of Biochemistry and Microbiology, University of Chemistry and Technology Prague, Prague 6 166 28, Czech Republic; Department of Science, Technology and Society, Scuola Universitaria IUSS, Istituto Universitario di Studio Superiori, Piazza della Vittoria 15, 27100 Pavia, Italy; Department of Clinical Genetics and Genomics, Sahlgrenska university hospital, Gothenburg, Sweden; Department of Laboratory Medicine, Institute for Biomedicine, Sahlgrenska Academy, University of Gothenburg, Gothenburg, Sweden; Institute of Clinical Medicine, Dept. Oncology and Pathology, Lund University, Sweden; Department of Clinical Research, Copenhagen University Hospital Amager and Hvidovre, Copenhagen, Denmark

**Keywords:** variant effects, cancer genomics, protein structures, free energy calculations, protein stability, protein function, long-range structural communication

## Abstract

The role of genomic variants in disease has expanded significantly with the advent of advanced sequencing techniques. The rapid increase in identified genomic variants has led to many variants being classified as Variants of Uncertain Significance or as having conflicting evidence, posing challenges for their interpretation and characterization. Additionally, current methods for predicting pathogenic variants often lack insights into the underlying molecular mechanisms. Here, we introduce MAVISp (<u>M</u>ulti-layered <u>A</u>ssessment of <u>V</u>arIants by <u>S</u>tructure for <u>p</u>roteins), a modular structural framework for variant effects, accompanied by a web server (https://services.healthtech.dtu.dk/services/MAVISp-1.0/) to enhance data accessibility, consultation, and reusability. MAVISp currently provides data over 1000 proteins, encompassing more than eight million variants. A team of biocurators regularly analyzes and updates protein entries using standardized workflows, incorporating free energy calculations or biomolecular simulations. We illustrate the utility of MAVISp through selected case studies. The framework facilitates the analysis of variant effects at the protein level and has the potential to advance the understanding and application of mutational data in disease research.

## Introduction

We are witnessing unprecedented advances in cancer genomics, sequencing^1^, structural biology^2^, and high-throughput multiplex-based assays^3,4^. While sequencing approaches can identify alterations in the genome, understanding the molecular mechanisms of these variants remains a challenge. Although many variants in human genes associated with disease are currently known, the identification of their effects on human health is lagging behind^5^. Substantial evidence, which is necessary to classify variants according to their effects, is often lacking or contradictory in nature. Consequently, Variants of Uncertain Significance (VUS) or variants found to have conflicting evidence are continuously identified and reported in variant databases^67–11^. VUS remain an outstanding problem which complicate diagnosis and lead to suboptimal diagnosis or choice of therapy ^12^.

At the same time, the bioinformatics community has developed various approaches for predicting the impact of variants on human health, many of which are benchmarked against or complemented by experimental data and cellular readouts^13–17^ In this context, experimental multiplex assays deliver good quality and high-throughput assessment of the effect of variants on different readouts and have effectively been used to aid clinical variant interpretation. ^18,19^ These computational and experimental approaches allow to classify variants for their potential pathogenic or benign effects, which are then reported in different repositories and compendia^7–10^. In fact, computational methods are currently considered supporting evidence for variant classification, according to recent revisions of the American College of Medical Genetics and Genomics/Association for Molecular Pathology (ACMG/AMP) variant classification guidelines^20^. Variant effect predictors (VEPs), methods designed to predict the effect of a mutation at the genome or protein level, have made considerable progress, as outlined in recent reviews^21–23^. VEPs have classically relied on sequence data and variants with known classifications.

Nonetheless, in recent years, the advent of AlphaFold2^2,24,25^ and other similar methodologies has enabled the prediction of accurate three-dimensional (3D) protein structures and complexes, often with a quality comparable to experiments. This, in turn, enabled the inclusion of information about protein structure in machine learning models, which are among the best-performing available VEPs^21^. A well-known example of this is AlphaMissense^26^, which is based on a deep learning model similar to AlphaFold2. Additionally, it simultaneously learns to perform structure prediction and trains an unsupervised protein language model, thereby incorporating structural information into the prediction. The latter was then fine-tuned for a variant classification task. Approaches based on protein language models (such as ESM-1b^27^ or, more recently, ESM-2^28^ and ESM-3^29^), which are unsupervised models of protein sequence, have also shown good performance when used in variant effect prediction tasks^29,30^. ESM-3^29^ already incorporates structural information into its training, through specialized tokens, whereas protein sequence models have been used in conjunction with structural information in various ways^31,32^. Even a model such as GEMME, which is an epistatic model entirely based on sequence conservation, has been supplemented with structural information as structure-derived features in ESCOTT^33^. Rhapsody-2 is a VEP that incorporates features derived from protein structure and dynamics within a machine learning framework^34^. Finally, the ability to perform long and accurate biomolecular simulations and robust physical models allows the exploration of conformational changes and protein dynamics across different timescales^35^.

In previous pilot projects, we explored structure-based methods to analyze the impact of variants in coding regions of cancer-related genes, focusing on their consequences on the protein product^36–38^. We propose that these methodologies could be widely applied to study disease-associated variants. When formalized and standardized, this approach can complement existing methods for predicting pathogenic variants, such as the aforementioned AlphaMissense^26^. Most available VEPs estimate the likelihood of damaging effects of variants, but do not provide evidence of variant effects in relation to specific altered protein functions at the cellular level. On the contrary, with this contribution, we aim to link the effects of variants to specific underlying molecular mechanisms^38^. A mechanistic understanding of variant effects can help the design of strategies in disease prevention, genetic counseling, clinical care, and treatment. Moreover, from a fundamental research perspective, mechanistic knowledge is also essential for designing and prioritizing experiments to investigate the underlying molecular causes of disease.

Considering this, we developed MAVISp (<u>M</u>ulti-layered Assessment of <u>V</u>arIants by <u>S</u>tructure for <u>p</u>roteins) to enable high-throughput variant analysis within standardized workflows. MAVISp integrates results from VEPs and structure-based predictions of variant effects on several protein properties. The data are accessible through a Streamlit-based website for consultation and download (https://services.healthtech.dtu.dk/services/MAVISp-1.0/). Additionally, we maintain a Gitbook resource with detailed reports for individual proteins (https://elelab.gitbook.io/mavisp/).

With this publication, we provide data on *in silico* saturation mutagenesis for all possible variants at each mutation site with structural coverage for 1096 proteins and over eight million variants. New data and updates of existing entries will be continuously released. Currently, we are capable of processing up to 20 new proteins weekly, which are deposited in a local version of the database. The public database is updated quarterly. Based on recent statistics (https://elelab.gitbook.io/mavisp/documentation/coverage-and-statistics), we anticipate providing 80-100 new proteins with each update, along with additional modules for existing entries. In this manuscript, we provide an overview of the methodology and show examples of data analysis and application.

## Results

### Overview of MAVISp and its database

MAVISp performs a set of independent predictions, each assessing the effect of a specific amino acid substitution on a different aspect of protein function and structural stability, starting from one or more protein structures. These independent predictions are executed by the so-called MAVISp modules (**Fig. 1a**). MAVISp can be applied to individual three-dimensional (3D) protein structures and their complexes (*simple mode*) or to an ensemble of structures generated through various approaches (*ensemble mode*). The framework is modular, allowing all the modules or only a selected subset to be applied, depending on the case study. Each module relies on Snakemake, Dask workflows, or Python scripts, all of which are supported by specific virtual environments. The modules are divided into two main categories: (i) modules to retrieve and select structures for analyses (shown in orange in **Fig. 1a**), (ii) modules to perform analyses related to variant assessment or annotations (shown in blue in **Fig. 1a**). Each module includes a strictly defined protocol for computational analysis that can be carried out either step by step or automatically embedded in more comprehensive pipelines (Methods). They are designed to ensure consistency across all the proteins under investigation and to enhance reproducibility and repeatability. Our prediction modules are also complemented by available experimental data or already available predictions that can be integrated in the MAVISp dataset, such as those for VEPs (shown in green in **Fig. 1a**). All the resources used in the MAVISp framework are reported in **Table S1**, some of which have been developed within this work.

**Fig. 1.**
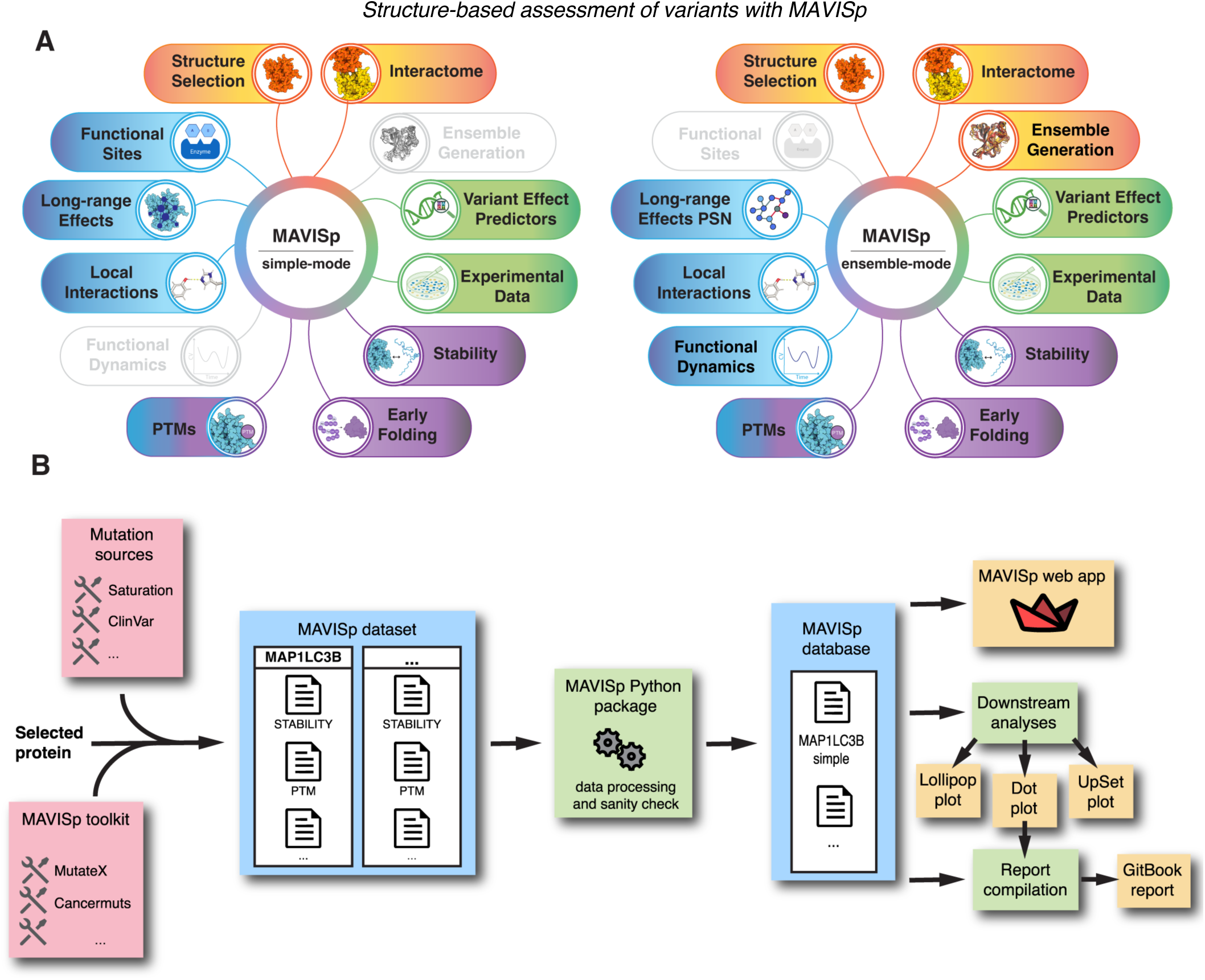
Overview of MAVISp components. (A) MAVISp includes different modules, each managed by workflow engines or dedicated tools. The modules highlighted in orange handle the selection and collection of protein structures, while the modules in blue and purple are dedicated to structural analyses of variant effects in relation to protein functional- or stability-related properties. Additionally, the framework provided modules with results from VEPs and scores derived by experiments, such as deep mutational scans (green). The procedure begins with a gene name, its UniProt and RefSeq identifiers and the desired structural coverage. For each gene, all the steps can be conducted on a standard server with 32-64 CPUs. The only exceptions are: i) the ENSEMBLE GENERATION module, which includes all-atom MD simulations, and ii) Rosetta-based calculations on binding free energies and folding/unfolding free energy calculations. Depending on the simulation length and system size, these might require access to HPC facilities. On the left, the *simple mode* for the assessment is illustrated, which uses single experimental structures or models from AlphaFold2 or AlphaFold3. On the right, the *ensemble mode* is schematized in which a conformational ensemble for the target protein or its complexes is applied. Hereby, we consider a conformational ensemble a collection of 3D conformations of the protein generated by a sampling method such as molecular dynamics or provided by NMR structures in the PDB (B) Scheme of the current workflow for the MAVISp database and websever. Biocurators apply specific workflows and protocols within each MAVISp module to generate structure-based predictions of changes linked to variants in each protein target. In doing so, they take advantage of the MAVISp toolkit as well as our mutation sources. The results are gathered into a text-based database. The data are further processed by the MAVISp Python package, which performs consistency checks, aggregate the data and outputs human-readable CSV table files, that make up the MAVISp database. These CSV files are imported by the Streamlit web app, powering the MAVISp webserver (https://services.healthtech.dtu.dk/services/MAVISp-1.0/), where the data are available for interactive visualization and download. In addition, the MAVISp database can be used to generate graphical representations of the data, such us dot plots, lollipop plots, and UpSet plots. Finally, based on the information gathered so far, we provide GitBook reports to facilitate the interpretation of the results: https://elelab.gitbook.io/mavisp/.

The modules are used in the context of the overall MAVISp workflow (**Fig. 1b**), which is designed to enable multiple biocurators to work concurrently and independently on distinct proteins. Data managers defined a priority list of targets that are analyzed in batches by biocurators, depending on the specific research project requirements. Additional targets of interest for the research community can be requested, as explained in the documentation on GitBook.

The workflow is designed as a set of consecutive steps that act on a protein of interest at a time. As the first step, once a protein of interest has been selected, a biocurator retrieves structural and functional information about it, along with key identifiers (e.g., gene name, UniProt AC, RefSeq identifier) for the next steps. Additionally, the biocurator proposes a trimming strategy for the protein, e.g., identifying one or more sets of contiguous residues in the protein structure that can effectively serve as input for the prediction steps. This step entails considering only well-structured and high-accuracy regions of our proteins, which is crucial since most MAVISp modules are not designed to handle large intrinsically disordered regions. In selected cases, to avoid potential bias in our structural calculations, the curator may edit the structure by removing long disordered inclusions in structured regions. Furthermore, in the MAVISp *ensemble mode*, where he ENSEMBLE GENERATION module should be carried out, the biocurator identifies the initial structures for the simulations to be performed on the protein target in its free or bound state with other biomolecules and performs the necessary simulations to obtain the final structural ensemble. Once the protein structure or structural ensemble, depending on the mode, is available, the biocurator works with each available module and obtains: i) a list of variants that MAVISp will annotate (see Materials and Methods for details) and ii) the final predictions for each module. To do so, biocurators adhere to strict workflows for data collection based on a set of procedures codified in each module, which is mostly automated via the use of Snakemake pipelines. Once this is completed, the MAVISp data managers will import and aggregate the data using the MAVISp Python package (https://github.com/ELELAB/MAVISp). This step also allows to perform sanity checks, per-module data classifications, and write the results in a human-readable table format, constituting the MAVISp database. The database files are the first product of MAVISp and contain the relevant collected data and metadata for each of the identified variants (https://services.healthtech.dtu.dk/services/MAVISp-1.0/).

The datasets from the MAVISp database can then be further used in two ways. First, biocurators or data managers can perform a set of analyses, referred to as downstream analyses, which are generated downstream of database creation. These analyses result in the generation of publication-ready figures that summarize the predicted effects for each variant and assist results interpretation.

Furthermore, the biocurators use data from the downstream analysis to create a report in GitBook (https://elelab.gitbook.io/mavisp/), using a standard Markdown template and a semi-automated procedure. Biocurators and data managers also act as reviewers for reports created by their peers. A review status is assigned to each GitBook entry to guide users regarding the quality and integrity of the curated data. To achieve this, we defined four review status levels (i.e., stars) for each protein entry (https://elelab.gitbook.io/mavisp/documen-tation/mavisp-review-status).

Finally, the MAVISp database is presented through a user-friendly Streamlit-based website (https://services.healthtech.dtu.dk/services/MAVISp-1.0/). The web app includes various visualizations to aid the interpretation of MAVISp results that are essentially equivalent to the downstream analyses outlined above: (a) a dot plot displaying classifications for each variant across MAVISp modules, experimental data (if available), and the VEP results, (b) a lollipop plot aggregating relevant mechanistic indicators (i.e., MAVISp-identified effects at the structural level) associated with potentially pathogenic variants, and (c) an interactive representation on the 3D structure, showing the localization of mutation sites identified in (b). These features are designed to support the interpretation of results and facilitate the identification of variants with specific mechanisms and multiple effects. The source code for the MAVISp Python package and the web application are available on GitHub (https://github.com/ELELAB/MAVISp), while the complete dataset can be downloaded from an OSF repository (https://osf.io/ufpzm/). The OSF repository also include previous version of the database. Both source code and data are freely available and released under open-source or free licenses.

We invite requests on targets or variants that are not yet available in MAVISp or scheduled for curation. We also welcome contributors as biocurators or developers, pending training and adherence to our guidelines (https://elelab.gitbook.io/mavisp). To facilitate entrance into the MAVISp community of biocurators and developers, we organize training events, research visits and workshops.

Notably, a comprehensive update will be conducted annually to incorporate new versions of external tools or resources used by MAVISp, ensuring that resources remain current. Moreover, we continuously expand our toolkit and develop new modules to enable even more comprehensive assessments. The criteria for including new methods and approaches in the framework are detailed in the GitBook documentation (https://elelab.gitbook.io/mavisp/documentation/how-to-contribute-as-a-developer).

### MAVISp modules for structure collection and selection

MAVISp includes various modules to select and model the structures of interest in both *ensemble* and *simple mode* (**Fig. 1a**). The STRUCTURE SELECTION module enables biocurators to identify the starting structure for their study, both for models of the free and bound states of the protein of interest. This module includes structure retrieval from the Protein Data Bank (PDB)^39^, the AlphaFold Protein Structure Database^25^, or through the generation of initial models with AlphaFold3^40^, AlphaFold2^2^ and, AlphaFold-multimer^24^. In addition, it streamlines the selection of structures in terms of structural quality, experimental resolution, missing residues, amino acidic substitutions with respect to the UniProt reference sequence, as well as the AlphaFold per-residue confidence score (pLDDT), integrating tools such as PDBminer^41^. Using AlphaFill^42^ further assists in identifying cofactors to be included in the model structure or to identify mutation sites that should be flagged, if located in the proximity of a missing cofactor in the structure model. When necessary, a workflow is available to reconstruct missing residues or design linkers to replace large, disordered loops within structured domains (Methods).

According to the protocol established for the generation of the models, we retain 3D structures with reasonable accuracy based on parameters such as pLDDT, Predicted Aligned Error (PAE), and pDOCKQ2^43^. In addition, the module includes protocols based on AlphaFold ^24,44^ or comparative modeling^45,46^ when the complex between the protein target and the interactor involves Short Linear Motifs (SLiMs).

The INTERACTOME module aids the identification of protein interactors for the target protein and their complex structures by querying the Mentha database^47^, the PDB, and experimentally validated proteome-wide AlphaFold models ^48^, as well as the STRING database^49^ (Methods). Once a suitable set of interactors has been identified, the information is used to predict protein complex structures, which are then utilized in the subsequent steps (i.e., the LOCAL_INTERACTIONS module, see below).

The ENSEMBLE GENERATION module allows the use of structural ensembles from different sources, such as NMR structures deposited in PDB, coarse-grained models for protein flexibility (e.g., CABS-flex^50^) or all-atom Molecular Dynamics (MD) simulations (with GROMACS^51^ and PLUMED^52,53^) of the protein structure or its complexes. The choice of the method to be used is based on the required accuracy of the generated ensemble and the available computational resources. Once individual structures or structural ensembles for the protein candidate are selected – either alone or with interactors - the analysis modules can be used.

### MAVISp modules for structural analysis

MAVISp integrates different analysis modules for both *ensemble* and *simple mode* (**Fig.1a**). The minimal set of data required to import a protein target and its variants into the MAVISp database includes the results from the STABILITY and PTM modules, along with predictions from VEPs. The STABILITY module is devoted to estimating the effects of the variants on the protein structural stability using folding free energy calculations (Methods). This module leverages workflows for high throughput *in silico* mutagenesis scans^54,55^ and a newly implemented protocol for RaSP^56^ (Methods). All the methods used in this module predict change of free energy of folding upon the insertion of an amino acid substitution, and predictions are performed using FoldX, Rosetta, or RaSP. Once these predictions have been collected, MAVISp applies a consensus approach to classify the effect of the variants (Methods). The defined thresholds for changes in free energy are based on evidence that shows that variants with changes in folding free energy below 3 kcal/mol do not exhibit a marked decrease in stability at the cellular level ^57,58^. Thus, MAVISp defines the following classes for changes in stability: stabilizing (ΔΔG ≤ - 3 kcal/mol with both methods, FoldX and Rosetta or RaSP), destabilizing (ΔΔG ≥ 3 kcal/mol), neutral (-2 < ΔΔG < 2 kcal/mol), and uncertain (-3 < ΔΔG ≤ -2 kcal/mol or 2 ≤ ΔΔG < 3 kcal/mol). A variant is also classified as uncertain if the two methods would classify the effect of the variant differently. Since March 2024, we adopted the consensus between RaSP and FoldX as a default for data collection, after performing a benchmark using the MAVISp datasets (**Supplementary Text S1** and https://github.com/ELELAB/MAVISp_RaSP_benchmark). RaSP provides a suitable solution for high-throughput data collection compared to the CPU-intensive scans based on Rosetta. In low-throughput studies, where we focus in detail on a target protein, we can include Rosetta data, which are computationally more demanding.

The LOCAL INTERACTION module can be applied if the STRUCTURE SELECTION and INTERACTOME modules identify at least a suitable structure of the complex between the target protein and another biomolecule. The LOCAL INTERACTION module is based on estimating of changes in binding free energy for variants at protein sites within 10 Å of the interaction interface, using protocols and consensus strategies that mirror those for STABILITY. In this case, we use a combination of FoldX and Rosetta calculations (Methods). Binding free energy thresholds are set based on the expected error margins of the predictors, approximately ±1 kcal/mol, as outlined by the authors of the methods and in accordance with general good practice in the literature. This approach addresses the scarcity of experimental datasets on amino acid substitutions that impacting protein-protein interactions^59–61^, which are often constrained by system heterogeneity, limited mutation numbers, or both, thereby complicating reliable benchmarking. We rely on a consensus approach between the results of FoldX and Rosetta on changes in binding free energies upon amino acid substitution. We classify a variant as stabilizing (both methods predict ΔΔG <= -1 kcal/mol), neutral (-1 kcal/mol < ΔΔG < 1 kcal/mol) or destabilizing (ΔΔG >=1 kcal/mol). Cases in which the two methods disagree on the classification, or for which we do not have a prediction for both methods, and the side chain relative solvent accessible area of the residue is >= 25%, are classified as uncertain. This is because, in high-throughput data collection, we cannot exclude the possibility that the site interacts if it is solvent exposed, as often in structural biology, only part of the 3D structures of protein-protein complexes are available or can be modelled. We also included support for LOCAL INTERACTION for protein and DNA interactions, as well as for homodimers. Notably, a strength of our approach is to provide annotations for the effects of protein variants on various biological interfaces for the same target protein.

In the *ensemble mode*, the STABILITY and LOCAL INTERACTION modules are used on ensembles of at least 20-25 structures from the simulations or on the three main representative structures upon clustering, depending on the free energy calculation scheme to apply. The results obtained for each structure are then averaged, and classification is performed with the same strategies we use in *simple mode* using these average values. This approach is used to mitigate limitations due to lack of backbone flexibility when these free energy methods are applied to just one single 3D structure^38,54,62,63^.

The LONG-RANGE module applies coarse-grained models to estimate allosteric free energy changes upon amino acid substitution based on AlloSigMA2^64^. The protocol followed by the LONG-RANGE module has recently been updated and benchmarked using experimental data from deep mutational scans^65^. Details on the parameters and steps for analysis are also provided in the Methods. Variants are annotated as destabilizing (positive changes in allosteric free energy), stabilizing (negative changes in allosteric free energy), mixed effects (both conditions occur), or neutral if the variant does not cause any significant change. Additionally, variants that do not cause a significant change in residue side-chain volume are annotated as uncertain. In the *ensemble mode*, we applied graph theory metrics based on changes in the shortest communication paths using atomic contact-based Protein Structure Network^66^. This analysis, combined with the AlloSigMA2 data, allows pinpointing variants with long-range effects to functional sites or protein pockets that could serve as interfaces to recruit interactors or ligands.

The FUNCTIONAL SITES module in *simple mode* allow to evaluate the effect of variants at (or in the proximity of) the active site of enzymes or cofactor binding sites of proteins and it is based on analyses of contacts with the second sphere of coordination of the residues belonging to these sites (see Methods).

The FUNCTIONAL DYNAMICS module in *ensemble mode* includes enhanced sampling simulations to further assess the local or long-range effects of a variant. As a first example, we applied this class of methods to validate the long-range effects predicted for p53 variants on the DNA-binding loops^38^, and included such results in the MAVISp database.

The PTM module currently supports phosphorylation only, annotating the effect of variants at phosphorylatable sites. It evaluates how the loss or changes of phosphorylation sites may impact protein regulation, stability, or interaction with partners. To this goal, the module collects analyses and annotations such as solvent accessibility of the mutation site, inclusion of the site in phosphorylatable linear motif, comparison between predicted changes in folding or binding free energy upon amino acid substitution or upon phosphorylation at the site of interest. In the module, we applied a custom decision logic (**Supplementary Text S2**) to derive the classification for each variant as neutral, damaging, unknown effect, potentially damaging or uncertain. The identification of the phosphorylation sites in the PTM module is based on known experimental phosphosites and SLiMs, as retrieved by Cancermuts^67^. These data are complemented by a manually curated selection of phospho-modulated SLiMs (https://github.com/ELELAB/MAVISp/blob/main/mavisp/data/phospho-SLiMs_09062023.csv). For solvent-inaccessible phosphorylatable residues, the effects are classified as uncertain in the *simple mode*. In these cases, the *ensemble mode* is required to investigate wheatear a cryptic phosphorylated site may become accessible upon conformational changes^68,69^. Of note, the current version of the PTM module has been designed based on fundamental principles on how phosphorylation can affect the protein structure and should be used to identify variants for further investigation, particularly for experimental research. Benchmarking the effectiveness of this module would be difficult at present time, given the relatively small number of amino acid substitutions that can affect phosphorylation currently present in the MAVISp database, especially considering those for which experimental data is available. To this purpose, we are currently in the process of curating and including more proteins relevant to benchmarking the PTM module. These will include experimental data on protein stability and protein-protein interactions upon phosphorylation ^70^ ^71^.

MAVISp includes further analyses and annotations, such as predictions on regions involved in early folding events^72^, pLDDT score, secondary structure, and side-chain solvent accessibility, which can assist in the interpretation of the results.

### Variant Effect Predictors included in MAVISp

MAVISp provides annotations for the variant interpretation reported in ClinVar^9^, or calculated with REVEL^73^, DeMaSk^74^, GEMME^14^, EVE (Evolutionary model of variant effect)^75^, and AlphaMissense^26^. In MAVISp, each of them is handled by a separate module. The results of these VEPs can be combined with the results from the MAVISp structure-based modules to understand variant effects and to prioritize variants for other studies, as detailed in the examples below.

### Sources of variants supported by MAVISp

By default, we apply in silico saturation mutagenesis, which means that we provide predicted effects for each variant of a target protein at any position that has a structural coverage. Additionally, all variants reported for the target protein in COSMIC, cBioPortal, and ClinVar are annotated within MAVISp. We routinely update and maintain the entries in the MAVISp database to include up-to-date annotations using Cancermuts^67^. All Cancermuts annotations for MAVISp and other protein targets are also available at the Cancermuts web-server, https://services.healthtech.dtu.dk/services/Cancermuts-1.0/. In addition, annotations from lists of variants from other studies, such as data on cohort-based or nationwide studies or other disease-related genomic initiatives, can be manually introduced.

Currently, MAVISp includes data on eight+ million variants from 1096 proteins (at the date of 20/11/2025). An overview of the currently available data and how to use them to address different research questions is described in detail in the next sections. The first targeted studies in which MAVISp has been applied to understand variants impact in rare genetic diseases^76^ or involved in cancer hallmarks^77,78^ are also suitable examples

### Interpretation of the results of MAVISp

MAVISp provides a comprehensive set of results for many variants; therefore, we have devised a few strategies that can be useful to make sense of the MAVISp data for a few common use cases that users might encounter.

One of the most important outputs from the downstream analyses, of MAVISp is the so-called dot plot, which is available on the GitBook reports or released within the target studies of specific proteins (see below for examples). A dotplot can also be generated within the MAVISp webserver in the “Classification” tab, for up to 50 variants of choice simultaneously. This plot showcases i) the classification of the different VEPs integrated in MAVISp, ii) the classification performed by each MAVISp module, iii) the classification of variants in ClinVar, when available, as variant label colors. The code to generate dot plots from MAVISp csv file is also available in GitHub (https://github.com/ELELAB/mavisp_accessory_tools/tree/main/tools). The MAVISp modules classification has a different meaning depending on the considered module: a variant classified as damaging for a VEP usually means it is predicted as functionally damaging or pathogenic (depending on the predictor), while a variant classified as damaging for stability just means that the variant is predicted to compromise the structural stability of the protein, and one classified as damaging by the long range module is predicted to have significant long-range effects, and so on. Another representation which depends on further processing of a text output created by dot_plot.py (i.e., alphamissense_out.csv) provides a concise representation of the classes of mechanistic indicators found for each variant in the form of lolliplots. Lolliplots are also available in the GitBook report or in the “Damaging mutation” tab on the website, that shows only those variants that are at the same time: i) classified as pathogenic for AlphaMissense, ii) classified as loss-of-fitness or gain-of-fitness by DeMaSk and iii) damaging for the respective structure-based module of MAVISp. The downstream analysis toolkit also provides the code to prepare upset plots or venn diagrams for the variant source (as reported in Gitbook).

Consulting the available dot plot for an entry of interest is therefore the most straightforward place to start to access MAVISp data. To identify a subset of variants of interest, we have defined the following strategy for a data-driven discovery of variants of interest with little other information (i.e. VUS, conflicting evidence or variants not reported in ClinVar). In this case, the dot plot allows to understand first which variants are predicted to be pathogenic, by using the AlphaMissense classification; these are the ones reported as Damaging in the AlphaMissense row. For these, we also consider the output of DeMaSk, that define whether the variant is classified as gain-of-fitness or loss-of-fitness. If a variant fullfil these criteria, we then consider the structure-based MAVISp predictions for mechanistic indicators, that give us one or more explanations of the reason for the effect of the variant. For instance, the variant could be destabilizing the protein structure and will be reported with an altered stability as mechanistic indicator. Another common use case is to use MAVISp to get a mechanistic interpretation of variants already known in ClinVar. In this case, if the variant already has an interpretation of Pathogenic, Likely pathogenic, Benign, or Likely benign, we can just refer to the MAVISp mechanistic interpretation.

Importantly, researchers should always refer to specific biological or phenotypical contexts when interpreting predictions from MAVISp, including their knowledge of the biological role the protein investigation has or concerning the nature of the disease of interest. For instance, predictions might lead to different conclusions if the protein under consideration is from a tumor suppressor or from an oncogene.

In the next section we illustrate some of the applications of data collected with MAVISp through case studies (**Table S2** for mapping of case studies and modules).

### COSMIC Tumor Suppressor Genes and Oncogenes

At first, we prioritized MAVISp data collection of known driver genes in cancer, i.e., tumor suppressors and oncogenes. To this goal, we collected data for the COSMIC Tumor Suppressor Genes (COSMIC v96), while the collection of the COSMIC Oncogene and Dual Role targets is ongoing. Furthermore, we have been including genes reported as a candidate driver by the Network of Cancer Genes (NGC)^79^.

The MAVISp datasets on cancer driver genes can assist the identification of molecular mechanisms of predicted or known pathogenic variants in these genes, as well as to aid the characterization of Variants of Uncertain Significance (VUS). A recent example is the study we performed on BRCA2^78^.In this study, we analyzed BRCA2 variants reported in ClinVar, comparing the predictions from the STABILITY and LOCAL INTERACTIONS modules of MAVISp with results from a multiplex assay which measured the impact of these variants on cell viability. We were able to explain the effect of 84 BRCA2 variants, which were classified as non-functional by the assay, and for which MAVISp predicted effects on protein stability or binding to the binding partner SEM1.

In the case of tumor suppressors, the identification of variants that might lead to loss of function is particularly important. Given structure-function relationship in proteins, structural stability represents a key determinant that can be disrupted by amino acid substitutions, potentially resulting in local or more drastic misfolding and loss of function^80^ As an example of loss of function due to changes in stability, we report the analysis of the MAVISp entry for the tumor suppressor BLM, a DNA helicase involved in DNA replication, recombination and repair^81^. We identified a total of 1170 predicted destabilizing variants according to the STABILITY module, of which 45 annotated in ClinVar (**Fig. 2a**). Among these, 82% destabilizing variants was found in structured regions of the protein, while the remaining 18% are located in disordered residue stretches (**Fig. 2b**). Of the ClinVar-reported variants, 42 are classified as VUS. Y811C and C901Y are reported with conflicting interpretations and only G952A is reported as likely pathogenic.

**Fig. 2.**
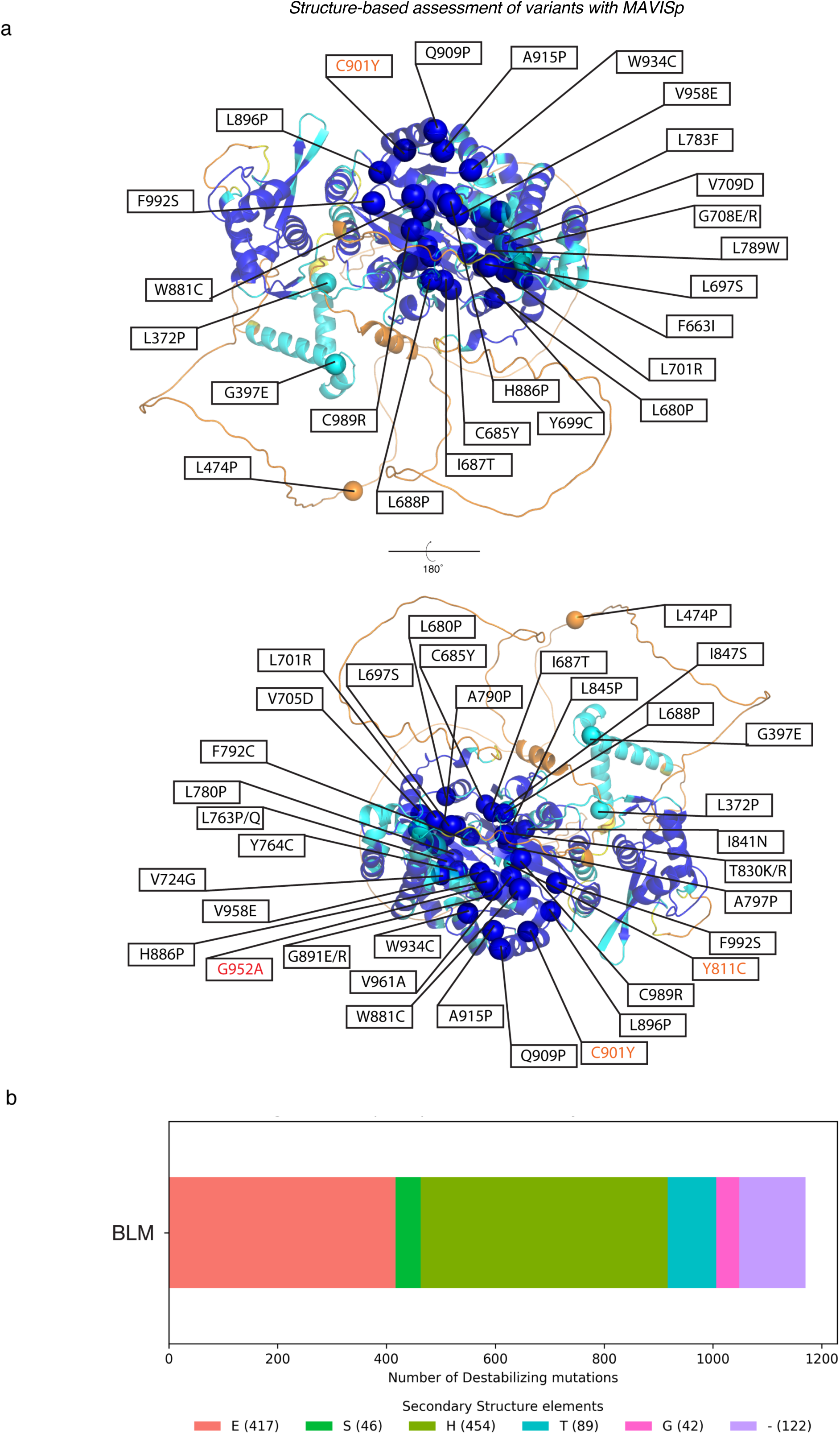
Variants with effects on structural stability in the tumor suppressor protein BLM. (a) The cartoon representation shows the trimmed mode BLM368-1290 and the spheres highlight the Cɑ atom of the 41 positions harboring 45 variants predicted as destabilizing by the MAVISp STABILITY module (RaSP/FoldX consensus) and annotated in ClinVar. Among these, Y764C, G891E, and L896P are also reported in CBioPortal, whereas F663I, L845P and C901Y are also reported in COSMIC. The two views correspond to the same domain rotated by 180°. The backbone and spheres are colored according to the AlphaFold pLDDT scores, i.e., blue - very high (pLDDT > 90), cyan - confident (70 < pLDDT <= 90), yellow - low (50 < pLDDT <= 70), and orange - very low (pLDDT <= 50). The labels indicate the mutation sites and the corresponding variants and are colored by ClinVar classification., uncertain significance (black), conflicting interpretation of pathogenicity (orange), and likely pathogenic (red). (b) The stacked bar plot shows the distribution of destabilizing BLM variants across secondary structure elements as defined by DSSP ((i.e., H = ɑ-helix, B = residue in isolated β-bridge, E = extended strand, participates in β ladder, G = 3-helix (310 helix), I = 5-helix (π-helix), T = hydrogen bonded turn, S = bend, and “-” = no secondary structure identified). The results refer to the data available in the MAVISp database on 12^th^ September 2025. More information about BLM analyses with MAVISp can be found in the corresponding GitBook report: https://elelab.gitbook.io/mavisp/proteins/blm

These results provide a starting point for variant characterization and prioritization. As suggested in the previous section, our results can be used to guide the selection of a subset of variants that have a predicted pathogenic impact from AlphaMissense, with a loss-of-fitness signature according to DeMaSk and that we predict damaging for stability., These would be suitable candidates for experimental validation. Concerning BLM, MAVISp identifies 41 ClinVar VUS or variants with conflicting evidence that could be prioritized according to these criteria (**Table S3**).

For example, depending on the size of the library to validate, methods such as flow cytometry sorting or cycloheximide chase assays^82,83^ or use approaches based on multiplex technologies^84–87^ would be useful to validate our predictions

### Integration of MAVISp data with experimental data

A useful feature of MAVISp is a dedicated module to curate and import experimentally derived scores on the effects of the variants on different biological readouts (i.e., the EXPERIMENTAL DATA module, **Fig. 1a**). These data can be directly compared with the structural properties we predict with MAVISp, for a variety of purposes. For example, they can serve as additional layer of information respect to the structure-based mechanistic indicators themselves. Additionally, as done in the aforementioned BRCA2 study, they can be used as a source of information for variants with a known detrimental effect that can depend on different mechanisms of action for each variant, which can be investigate using MAVISp. In cases such as this, MAVISp helps identifying the possible mechanism for which variants have an effect, for further in-depth investigation.

Experimental data can also be used to validate the results of certain MAVISp modules, for cases in which the predicted structural properties are related to the experimentally tested biological readouts. Deep mutational scans can also be used to benchmark or tune the thresholds used for classification performed by the MAVISp modules, including structural properties. In this context, the format of MAVISp database files is handy for further data processing, for example using biostatistical models or machine learning. In the case of PTEN, we included data from available deep mutational scans, reporting on the effect of mutations on cellular abundance or phosphatase activity^84,88,89^, in its MAVISp entry. Cellular abundance represents a critical property that is often perturbed by missense mutations, and that can be altered by changes in protein structural stability. We therefore compared predictions from the MAVISp STABILITY module—based on a consensus of RaSP and FoldX —with protein abundance scores obtained from VAMP-seq assays ^84,89,90^. To compare the classification obtained by the stability module with the experimental data, we considered how the abundance score from the experiment have been classified. Multiple classification strategies have been used for these data: on one side, the ProteinGym benchmark dataset^91^ applies a threshold based on the median of the abundance score (i.e., 0.77). Variants that scored lower than this threshold (>=22% reduction of abundance relative to the wild-type) were classified as low abundant, whereas those that scored higher were considered to be similar to the wild-type. The second classification followed the original PTEN study deposited in MaveDB (MaveDB ID urn:mavedb:00000013-a-1), which defines four abundance classes. In this scheme, the 5% lowest-abundance synonymous variants corresponded to a score of 0.71^84^, and variants were classified into bin 1 (low-abundant, both score and confidence interval < 0.71), bin 4 (WT-like abundant, both scores > 0.71), bin 2 (likely low-abundant, score < 0.71 but confidence interval > 0.71), and bin 3 (likely WT-like abundant, score > 0.71 but confidence interval < 0.71). For this analysis, we retained only variants in bins 1 and 4, to ensure an unambiguous classification. After applying these filters and excluding uncertain variants defined by the STABILITY module, the MaveDB-based classification contained 1690 variants. To enable a direct comparison between the two classification strategies, the ProteinGym-based dataset, which initially comprised 3211 variants, was filtered to include the same 1690 variants as the filtered MaveDB dataset. The two classification schemes were found to be largely concordant, differing only for variants with abundance scores between 0.71 and 0.77, which were considered damaging by ProteinGym and neutral by MaveDB.

**Fig.3 and Table 1** illustrate the performance of the MAVISp STABILITY classification against the classification of experimental data on protein abundance for PTEN. In this first comparison, we applied the same threshold suggested for this dataset from the benchmarking dataset ProteinGym^91^, which is based on the median value of the DMS scores. The consensus approach provided by the STABILITY module of MAVISp (accuracy 0.814) has an overall better performance in identifying variants that are found to be damaging in the assay than those predicted to cause damaging effects according to GEMME or DeMaSk (**Fig. 3b**). Nevertheless, this approach has a lower sensitivity (0.66) compared to GEMME. We thus wondered if the relatively low sensitivity we obtained was due to cases with experimental scores too close to the median (**Fig. 3c**). Additionally, in the original study for PTEN and as deposited in the MaveDB^92,93^, a different classification for the variant scoring based on four abundance levels was proposed, as detailed above. We thus performed a comparison of the MAVISp results with the experimental dataset for the PTEN experiment from MaveDB using the abundance level classes as a threshold (**Fig. 3c)**, resulting in increases sensitivity for the methods applied within MAVISp. The results on PTEN from MAVISp fits nicely with recent computational studies of PTEN variants using Rosetta calculations of protein stability and analyses of sequence conservation^16,17^.

**Fig. 3.**
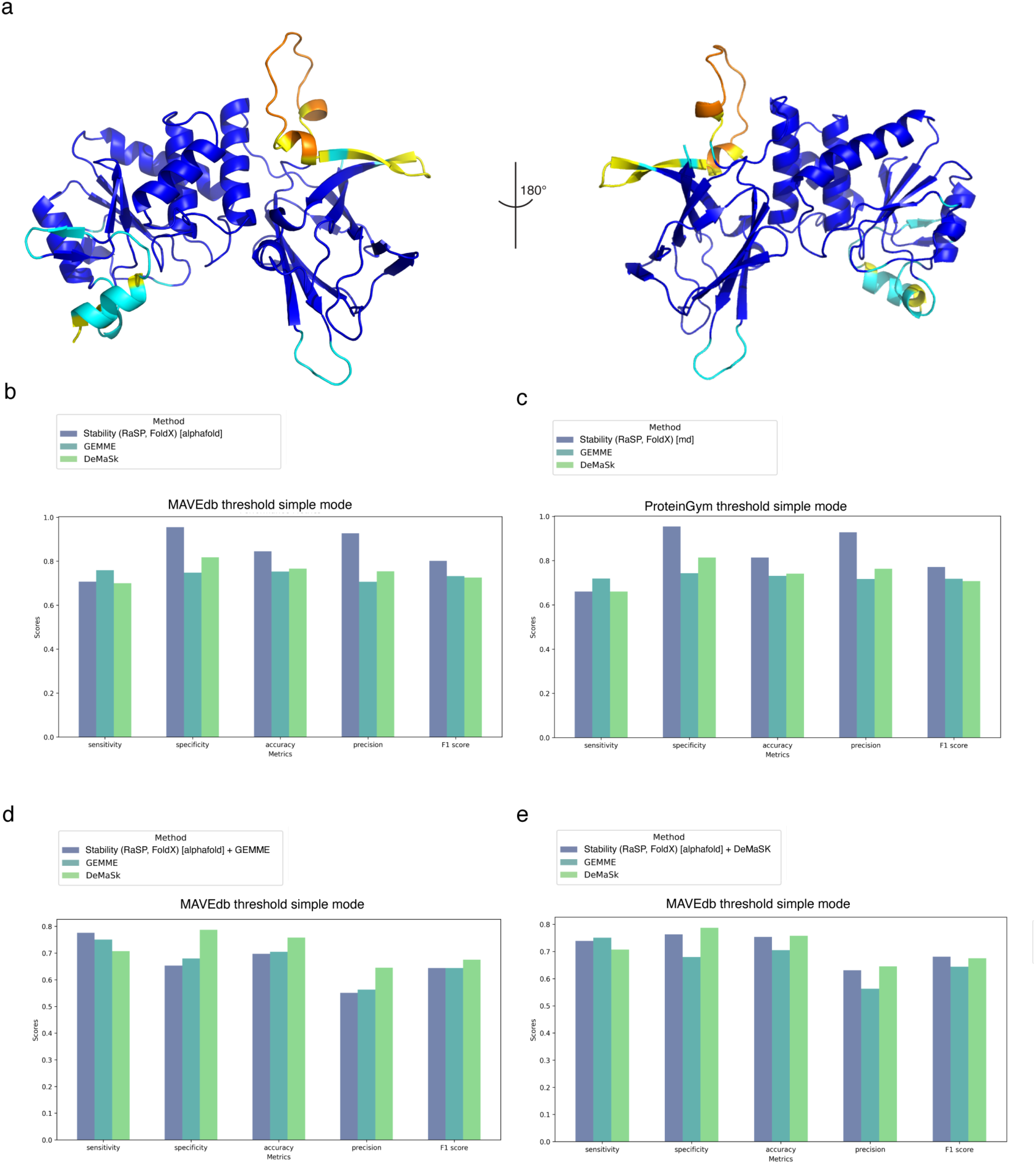
Comparison of GEMME, DeMaSk, and MAVISp STABILITY module predictions with experimentally-derived scores for protein abundance and phosphatase activity of PTEN. _(a) The trimmed AlphaFold structure (residues 1-351) of PTEN used for MAVISp stability module calculations is shown as a cartoon, colored according to pLDDT scores. (b-e) Histograms with performances of MAVISp STABILITY module, DeMaSk, and GEMME in predicting the effect of variants using VAMP-seq scores with ProteinGym (b) and MaveDB (c) thresholds. (d-e) illustrates the performances of the same tools or their combination against an experimental functional readout that assess the phosphatase activity at the cellular level._

**Table 1.** Performances of MAVISp modules and VEP predictors against experimental measurements of protein abundance and phosphatase activity for PTEN.

| Assay | column comparison | threshold_mode | sensitivity | specificity | accuracy | precision | F1 score |
| --- | --- | --- | --- | --- | --- | --- | --- |
| protein abundance assay | RaSP/FoldX consensus | ProteinGym | 0,666 | 0,954 | 0,814 | 0,928 | 0,771 |
|  | GEMME |  | 0,719 | 0,743 | 0,731 | 0,717 | 0,718 |
|  | DeMaSk |  | 0,66 | 0,814 | 0,741 | 0,763 | 0,708 |
|  | RaSP/FoldX consensus | MaveDB | 0,707 | 0,955 | 0,845 | 0,927 | 0,802 |
|  | GEMME |  | 0,759 | 0,748 | 0,753 | 0,706 | 0,732 |
|  | DeMaSk |  | 0,7 | 0,818 | 0,766 | 0,754 | 0,726 |
| phosphatase assay | RaSP/FoldX + GEMME | MaveDB | 0,776 | 0,653 | 0,697 | 0,551 | 0,644 |
|  | RaSP/FoldX + DeMaSk |  | 0,739 | 0,763 | 0,754 | 0,631 | 0,681 |
|  | GEMME |  | 0,751 | 0,68 | 0,705 | 0,563 | 0,644 |
|  | DeMaSk |  | 0,707 | 0,787 | 0,758 | 0,645 | 0,675 |

We next assessed whether MAVISp could also inform predictions of variant effects on PTEN phosphatase activity. Experimental data from a cellular phosphatase assay (MaveDB ID urn:mavedb:00000054-a-1) were classified as reduced (< 0.89), wildtype-like (0.89–1), or hyperactive (> 1). Here, we investigated whether integrating MAVISp STABILITY data with VEP results could enhance predictive power, since reduced stability is not the only possible mechanism for loss of phosphatase activity in mutated variants. We combined the STABILITY results with GEMME or DeMaSk, applying a priority logic in which damaging calls from GEMME/DeMaSk were given priority. This strategy produced performance comparable to GEMME or DeMaSk alone (**Fig.3d-e, Table 1**). Notably, combining changes in folding free energies with GEMME increased sensitivity (0.78) but reduced specificity (0.653), yielding an F1 score comparable to that of GEMME but lower than the one of DeMaSk (0.64; **Fig.3d-e, Table 1**).

Overall, with the examples in this section, we illustrate examples on how to use MAVISp data to compare predictions and experiments, as well as how to integrate MAVISp modules on structural properties with VEP results.

### Proteins involved in cancer hallmarks

To expand the contents of the MAVISp database, we have also been focusing on protein targets related to cancer hallmarks^94^, and in particular on proteins involved in cancer hallmarks related to protein clearance at the cellular level, i.e. the ability to escape cell death through apoptosis and autophagy, as well as kinases or transcription factors involved in the regulation of cellular proliferation. Mitochondrial apoptosis is tightly regulated by a network of protein-protein interactions between pro-survival and pro-apoptotic proteins. We investigated the mutational landscape in cancer of this group of proteins in a previous study^95^, which includes structural analyses with different *simple mode* MAVISp modules for both the pro-survival proteins BCL2, BCL2L1, BCL2L2, BCL2L10, MCL1 and BCL2A1, as well as the pro-apoptotic members of the family BOK, BAX and BAK1. In these analyses, the C-terminal transmembrane helix has been removed since the current version of our approach does not support transmembrane proteins or domains, illustrating an example on how the STRUCTURE SELECTION module works.

Autophagy is a clearance mechanism with a dual role in cancer. The autophagy pathway relies on approximately 40 proteins, constituting the core autophagy machinery^96^. As an example of the application of MAVISp to this group of proteins, we applied the *simple mode* to the markers of autophagosome formation MAP1LC3B and the central kinase ULK1, building on the knowledge provided by previous work^36,37^.

In the case of ULK1, we expanded our analysis to cover a larger part of the structure of the protein, meaning that both the N-terminal (residues 7-279) and C-terminal domains (837-1046) have been used for the MAVISp assessment. ULK1 also serves as an example of how to customize the trimming of an AlphaFold model to exclude disordered regions or linkers with residues featuring low pLDDT scores, in *simple mode*. In fact, their inclusion could lead to predictions of questionable quality. Disordered regions cannot be properly represented by a single conformation, and the *ensemble mode* would be necessary to derive more reliable conclusions. ULK1 featured 215 variants reported in COSMIC, cBioPortal and/or ClinVar, as shown in its dot plot (**Fig. 4A**), which was generated using the downstream analysis tools of MAVISp. Using the *simple mode*, 59 variants had predicted long-range mixed effects. Furthermore, eight had a damaging effect on stability, one had a damaging PTM effect on regulation (S954N), one had a possible damaging PTM effect in function (S1042T), and four variants (L53P, G183V, E191G, and L215P) are characterized by both effects on stability and long-range communication (**Fig. 4B-D**). Most of the variants that were predicted to have long-range and structure-destabilizing effects are in the N-terminal kinase domain of the protein, suggesting that mutations in this domain could result in the inactivation of ULK1 by compromising its 3D architecture. We then performed a one-microsecond MD simulation of the ULK1 N-terminal kinase domain (residues 3-279, PDB ID: 5CI7) to generate a structural ensemble for the MAVISp *ensemble mode*. In this case, we used an approach based on graph analysis from a contact-based PSN (Methods), as provided by the LONG-RANGE module, which verified if long-range communication occurs between mutation and response sites predicted by the coarse grain model used in the *simple mode*. The *ensemble mode* also validates the prediction on the effect of variants on stability that were done in *simple mode*, as it compensates for the none or limited mobility of the protein main chain that characterize the used in the STABILITY module. Overall, the application of the *ensemble* mode allowed to validate five variants with predicted long-range damaging effects (H72D, H72N, E73D, E73K, and R160L) and two variants with a damaging effect on stability (G183V and L215P). The predicted destabilizing (**Fig. 4E**) variant L215P has been also identified in samples from The Cancer Genome Atlas (TCGA) ^36^.

**Fig. 4.**
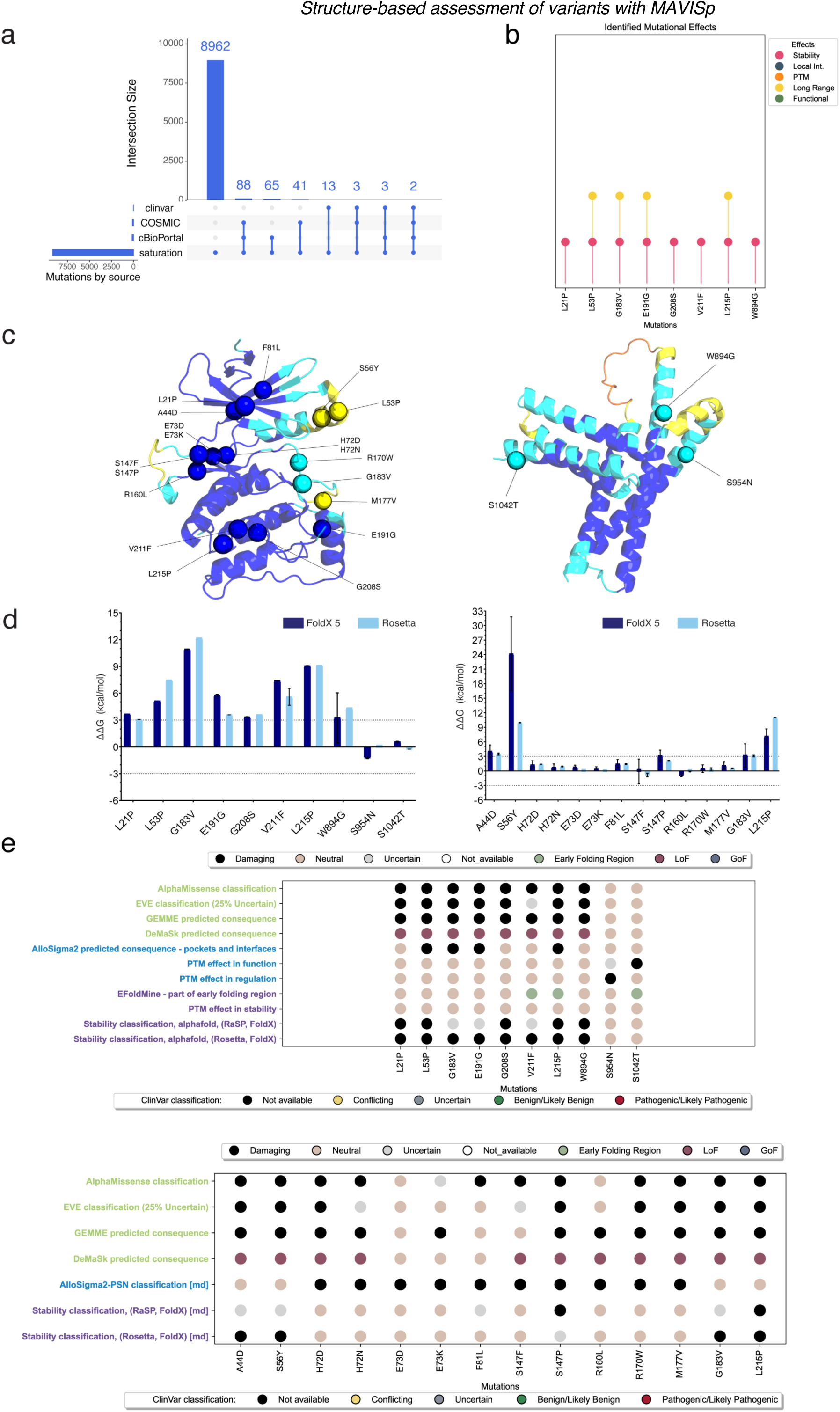
MAVISp *ensemble mode* to identify damaging variants in the autophagy kinase ULK1. a) We examined the central autophagy kinase ULK1 using MAVISp, generating a saturation of all possible variants within the N-terminal (residues 7-279) and C-terminal domains (residues 837-1046), leading to a total of 8,962 variants. Of these, 215 variants have been identified in COSMIC, cBioPortal, and/or ClinVar databases. b) Among the ones reported in the previous databases, eight variants were reported as pathogenic by AlphaMissense (L21P, L53P, G183V, E191G, G208S, V211F, L215P, and W894G) and among these, four variants are predicted to have a damaging effect on both protein stability and long-range communication (L53P, G183V, E191G, and L215P). c) Using MAVISp *simple* and *ensemble* modes, we identified 22 variants with destabilizing effects in terms of folding free energy, long-range effects, or PTM effects in regulation or in function. The mutation sites are highlighted with spheres on the AlphaFold models of the ULK1 N-terminal (left) and C-terminal (right) domains. d) We showed the predicted changes in folding free energy upon amino acid substitution for each of the 22 variants as calculated by the STABILITY module of MAVISp with MutateX and RosettaDDGPrediction with the *simple mode* (left) or with the *ensemble mode* (right). Interestingly, most of the variants that alter structural stability are located in the catalytic domain of the enzyme. This suggests potential mechanisms for ULK1 inactivation. e) Summary of the predicted effects on the 22 variants of ULK1 that have been found damaging with at least one MAVISp module with the *simple mode* (upper) or with the *ensemble mode* (lower) using the dot plot representation provided by the MAVISp toolkit for downstream analyses. Of note, the lower legend refers to the color of variants on the X-axis which are related to the ClinVar effect category.

The MAVISp entry of the autophagy marker MAP1LC3B provides an example on how the data for the LOCAL INTERACTION module can be obtained in a case of a protein that interacts with a functional motif embedded in intrinsically disordered proteins, i.e., a short linear motif (SLiM). MAP1LC3B in fact is able to bind to proteins harboring a so called LC3-interacting region (LIR)^97^. In MAVISp, we report the results for the effect on binding affinity of variants in MAP1LC3B or in its binding partners using three examples of this mode of interaction modeling the binding of MAP1LC3B with the LIR regions of its binding partner SQSTM1 (**Fig.5a**), ATG13, and Optineurin. In this case, we first applied the protocols for (phospho)-SLiM identification developed within the MAVISp framework (Methods) and PDBminer to identify possible starting structures. In the case of optineurin, we further model the flanking regions^77^. We identified ten variants annotated in ClinVar: nine reported as VUS (E102K, H86D, T29I, V91I, P2R, L44P, L44F, D56G, and R11L) and one as benign, i.e., E25Q (**Fig.5a**). MAVISp managed to predict a putative mechanistic explanation for the effect of four variants (**Fig.5b-d**): T29I is predicted to disrupt regulation by phosphorylation, L44P has an effect on both structural stability and long-range effects to distal sites, L44F and R11L have long-range effects (**Fig 5b**). Additionally, a variant found in cancer studies, P32Q, is predicted to have a detrimental effect on structural stability, confirming previous experimental results which showed propensity for aggregation^37^. Of note, this variant is identified with an uncertain prediction for the effect on stability in MAVISp simple mode, whereas two different approaches for generating a conformational ensembles accounting for protein dynamics predicted a destabilizing effect (**Fig 5d**). Additionally, all the variants with a mechanistic indicator from MAVISp are also predicted as pathogenic by AlphaMissense (**Fig.5c-d**) and are good candidates to further experimental studies for their effects on the autophagy flux or other functional readouts. V91I is likely to be benign variants since all the predictors identified neutral effects (**Fig. 5c-d**).

**Fig. 5.**
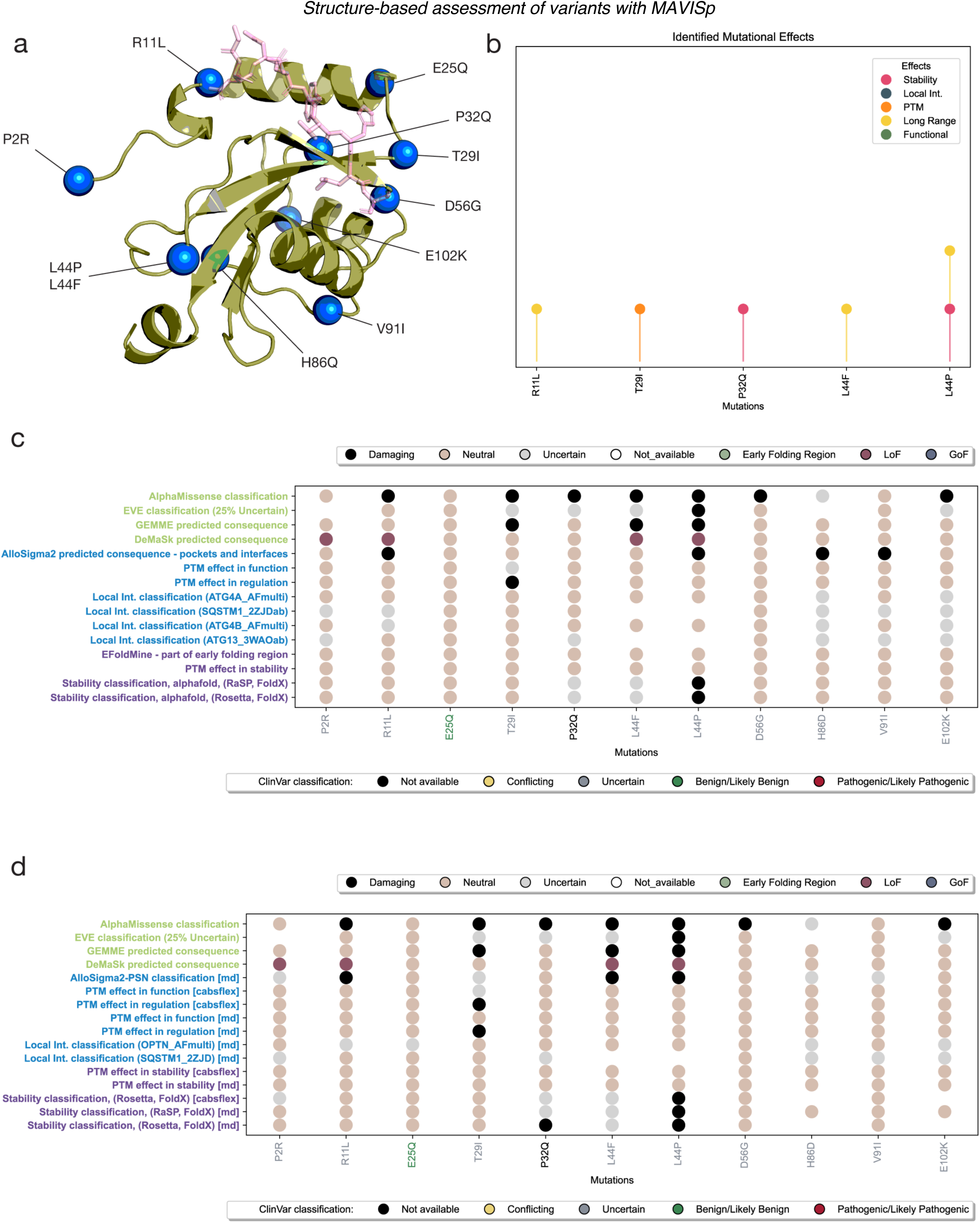
Analysis of MAP1LC3B VUS Variants from ClinVar. a) A structural model (PDB ID: 2ZJD) of the MAP1LC3B (green) interaction with the LIR motif of SQSTM1(pink) highlights ten ClinVar-reported variants (E102K, H86D, T29I, V91I, P2R, L44P, L44F, D56G, R11L and E25Q) along with the cancer-related variant P32Q. These variants are depicted as blue spheres on the structure. (b) Among these variants, five (R11L, T29I, P32Q, L44F and L44P) are predicted as damaging by AlphaMissense. Interestingly, L44P shows a predicted damaging effect on both long-range communication and stability. (c-d) Summary of the predicted effects on the 11 variants of MAP1LC3B as reported by MAVISp dot plot with the simple mode (c) or with the ensemble mode (d) using the dot plot representation provided by the MAVISp toolkit for downstream analyses. Of note, the lower legend refers to the color of variants on the X-axis which are related to the ClinVar effect category.

### Application of MAVISp to transmembrane proteins and to variants associated to other diseases

The STABILITY and LOCAL INTERACTION modules do not support predictions for variants in transmembrane regions. A survey on methods to predict folding free energy changes induced by amin on transmembrane proteins suggested that existing protocols, based on FoldX or Rosetta, are suitable for soluble proteins^98^. Therefore, the protocols implemented in the MAVISp modules for transmembrane proteins only retain those variants that are not in contact with the membrane. An example of a MAVISp entry for this class of proteins is PILRA, which has a low pLDDT score in the transmembrane region, and has been therefore excluded from the model, focusing on the analyses on the variants in the 32-153 region. In addition, we included other transmembrane proteins in the database such as ATG9A and EGFR.

PILRA is a protein target connected to neurodegenerative diseases^99^, along with KIF5A, CFAP410, and CYP2R1, illustrating the broad applicability of MAVISp to proteins involved in different diseases. Proteins associated with other diseases, such as TTR, SOD1, and SMPD1, have also been included in the MAVISp database. SMPD1 has been recently investigated in a targeted study using the *ensemble mode* of MAVISp together with other methodologies, validating our results by means of experimental data measuring the residual catalytic activity of enzyme variants^76^. As previously stated, MAVISp integrates curated experimental data for specific target proteins, which can be analyzed together with the results from the computational modules. To this goal, the dot plot representation provided by the downstream analyses toolkit of MAVISp and by the MAVISp database achieves a complete overview of both the experimental and the computational results (**Fig. 6a-c)** for SMPD1. Additionally, when a set of experimental data is available, it is possible to evaluate the correlation between predictions and experimental data (**Fig. 6d-e**). For SMPD1, we have obtained data on the residual catalytic activity of the enzyme for 135 variants^76^, available in the literature. Thanks to the MAVISp protocol, we predicted the effect of amino acid substitutions on changes in folding free energies as well as data for predicted functional effects from VEPs, which can be compared with the experimental data. The score values produced by the VEPs were mildly correlated with the residual activity measurements (Pearson correlation coefficient ∼0.6). Of note, most of the variants that have a predicted destabilizing effects on the stability are found at values of experimental residual activity lower than 20%, confirming what observed in our previous study^76^ and suggesting that changes in stability for SMPD1 can be help identifying damaging variants of this enzyme. Nonetheless, in this case, the experimental readout cannot be explained by stability changes alone. Thus, variants found with low residual activity and functionally damaging (GEMME and DeMaSk scores lower than -3 and -0.25, respectively) and that are neutral for stability according to MAVISp are good candidates for further investigation. For example, biomolecular simulations or computational chemistry methods could be used to investigate the effects of these variants on the catalytic mechanism of the enzyme and its lipid transport. Finally, variants, such as Y500H, which have a low residual activity, high loss-of-fitness scores and are uncertain for the STABILITY module, can be analyzed for their propensity to fall in early folding regions (see entry in the MAVISp database) and could be investigated in the *ensemble mode* using enhanced sampling simulations to accurately estimate their folding free energy profiles.

**Fig. 6.**
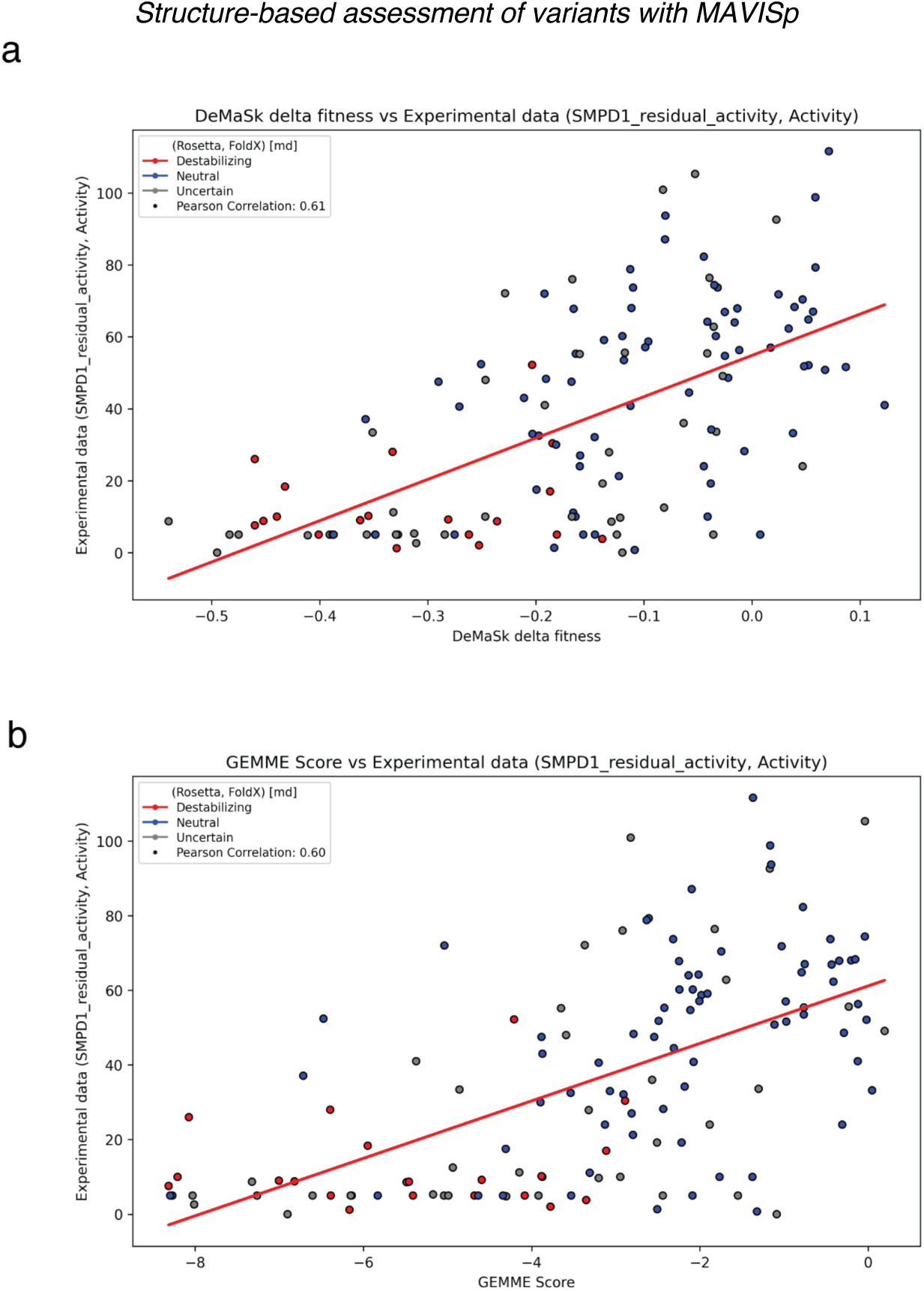
MAVISp, GEMME, and DeMaSk predictions on the impact of SMPD1 variant subset. A subset of SMPD1 variants, for which experimental data on enzyme activity have been selected, is shown with predictions from MAVISp, GEMME, and DeMaSk **a-b)** Scatter plots comparing DeMaSk (left) and GEMME (right) scores against experimental assay scores for enzymatic activity. The red line represents the regression, while the dotted line marks the threshold below which enzyme activity is considered inactive. Dots are colored based on the MAVISp STABILITY module classification (Rosetta/FoldX consensus): Destabilizing, Neutral, or Uncertain.

## Conclusions and Future Perspective

MAVISp provides a multi-layered assessment of the effects of variants found in cancer studies or other diseases using structural methods. MAVISp results are especially useful for variant interpretation and prioritization. These results can be useful as a complementary resource to available pathogenic scores or high-throughput experiments. MAVISp can help to pinpoint the effects linked to a pathogenic variant for further studies.

A significant advantage of MAVISp is its comprehensive coverage, expanding beyond clinically identified variants, by including novel variants yet to be characterized in other databases. This makes MAVISp a valuable resource for researchers and clinicians, facilitating the exploration of novel variants and their underlying pathogenic mechanisms. MAVISp can help on one side to associated mechanistic indicators to variants that are known or predicted pathogenic, as well as to aid in the characterization of the effects of VUS or variants with conflict evidence at the molecular level. Finally, we envision that MAVISp could become, in time, a community-driven effort and serve as a repository of data for the effects of disease-related variants more broadly. The results reported in MAVISp will provide an atlas of functional annotations for disease-related variants.

We have previously framed MAVISp in the context of others computational frameworks that collect data from different sources or integrate different structure-based methods to characterize variants^100^. This idea has led to the production of different attempts. Missense3D^101^ predicts the impact of variants on an array of structural features; ADDRESS^102^ includes predictions on stability and intermolecular contacts for variant found in UniProt humsavar; MUTATIONEXPLORER^103^ uses Rosetta and RaSP to predict the effect of amino acid substitutions on stability or binding, on user-provided structures; VUStruct^104^ selects relevant protein structures for the protein of interest and performs a wide array of predictions, including on the effect of variants on stability, binding surface, PTMs. The Genomics 2 Proteins^105^ portal includes data from several sources, including some overlapping with MAVISp such as Phosphosite or MaveDB, as well as features calculated on the protein structure. ProtVar^106^ also aggregates variant from different sources and includes both variant effect predictors, prediction of change on stability upon amino acid substitution, as well as prediction of complex structures. MAVISp, is, to our knowledge, the first resource to integrate data on binding free-energies, data derived from molecular dynamics simulations, as well as experimental data from different sources, and the first to integrate predictions on long-range effects as a database. While MAVISp has a lower coverage than others, it includes carefully curated manual steps, such as during protein structure preparation and simulation.

As the database grows, it will provide high quality data on different structural properties that can also be used for benchmarking purposes or as features in machine learning models. To this goal, the stringent data collection that we designed and present here is pivotal to build meaningful and accurate predictive models.

We would like to highlight previous studies that have demonstrated the usefulness of MAVISp and its protocols. For example, we have showcased the versatility of MAVISp in characterizing the effects induced by a redox post-translational modification of Cysteine (*S*-nitrosylation) using structural methods^107^. We focused on variants found in cancer samples for their capability to alter the propensity of cysteine to be *S-*nitrosylated, or a population-shift mechanism induced by the PTM. The collection of data using MAVISp modules has been pivotal to aggregate variants for each target of interest in the study on *S-*nitrosylation. The pipelines developed in the study of *S*-nitrosylation will be integrated within the MAVISp PTM module, extending it beyond support for phosphorylation, which is currently supported by MAVISp.

Alterations in transcription factors are often linked to aberrant gene expression, including processes such as proliferation, cell death, and other cancer hallmarks^108^. Different mechanisms are at the base of alterations in the activity of transcription factors in cancer, including point mutations. A previous study on TP53 served as a platform to develop different modules currently available in MAVISp ^38^. We thus aim to expand the MAVISp database to include more transcription factors. To this goal, one of the datasets under data collection covers the protein targets from the TRRUST2 database^109^, which includes experimentally characterized transcription factors and their targets, of which 150 have been already processed and included in the MAVISp database.

Furthermore, MAVISp provides pre-calculated values of changes in folding or binding free energies and other metrics that can also be reanalyzed in the context of other research projects. With the examples on PTEN and SMPD1 provided here, we introduced the curation of experimental data in MAVISp, as a source of experimental validation. The implementation of additional modules for MAVISp (e.g., degron^110^ and aggregation propensity^111^) would likely improve coverage of the diverse mechanisms regulating protein abundance. Of note, MAVISp supports either data from multiplex assays of variant effects or experimental data from literature mining of the biocurators. The purpose of collecting experimental data is to validate our findings, update protocols, and continuously improve the included methodologies. The reliability of our predictions depends on their alignment with experimental results, which can be used as reference data to benchmark and improve our predictions over time. The database currently includes 16 protein entries with experimental data.

At this stage, MAVISp can provide annotations for variants of transmembrane proteins exclusively in regions that are not in contact with the membrane. Recently published approaches^112^ could enable the application of the STABILITY module to transmembrane regions as well. In addition, we will include support to intrinsically disordered regions in the ensemble mode, designing new modules to reflect the most important properties of these regions.

We foresee that MAVISp will provide a large amount of data on structure-based properties related to the changes that can exert at the protein level, which could be exploited for design of experimental biological readouts, also towards machine-learning applications for variant assessment and classification or to understand the importance of specific variants in connection with clinical variables, such as drug resistance, risk of relapse and more.

## Methods

### Initial structures for MAVISp and STRUCTURE_SELECTION module

As a default, in the high-throughput data collection, we use models from the AlphaFold2 database^25^ for most of the target proteins and trim them to remove regions with pLDDT scores < 70 at the N- or C-termini or very long disordered linkers between folded domains. For proteins coordinating cofactors, in the low-throughput targeted studies, we re-modeled the relevant cofactors upon analyses with AlphaFill^42^ and where needed through MODELLER^113^. A summary of the initial structures used for each protein included in the database is reported in OSF (https://osf.io/y3p2x/). In selected cases, we have replaced long disordered loops with short residue stretches using a custom pipeline based on MODELLER (https://github.com/ELELAB/MAVISp_loop_replacer). This was done to avoid potential bias in our structural calculations, due to the arbitrary conformation of such loops and their spurious contacts with the rest of the structure. In addition, for proteins with transmembrane regions, we used the PPM (Positioning of Proteins in Membrane) server 3.0 from OPM (Orientations of Proteins in Membrane)^114,115^. For target proteins larger than 2700 residues, whose structures are not provided by the AlphaFold2 database, we model them using AlphaFold3.

The advantage of using AlphaFold-predicted structures in the default high-throughput data collection of MAVISp lies in their ability to achieve quality comparable to experimental data, as demonstrated in previous work^2^, and at the same time circumventing limitations typically associated with experimental approaches, such as artifacts, missing atoms, and incomplete or absent residues.

### INTERACTOME module

In the INTERACTOME module, implemented in the freely available PPI2PDB toolkit (https://github.com/ELE-LAB/PPI2PDB), we identify known interactors of the target protein by extracting data from the Mentha database^47^ and match them to available PDB structures, using the *mentha2pdb* script. Mentha2pdb also examines experimentally validated dimeric complexes generated with AlphaFold2 from the HuRI and HuMAP databases by Burke et al.^48^. Mentha2PDB provides annotations of the interactors and generates input files for AlphaFold-Multimer.

Complementarily, we retrieve interactors from the STRING database^49^ and process them analogously using our STRING2PDB tool, which maps STRING interactions to available PDB structures. The tool restricts retrieval from the physical subnetwork of STRING with evidence of interaction supported by either curated database annotation or experimental data.

As a final step, we aggregate all interaction data for the target protein into a single table, ranking interactors primarily by Mentha and secondarily by STRING score to prioritize experimentally supported pairs. We then add complexes retrieved directly from the PDB via pdbminer-complexes (https://github.com/ELELAB/MA-VISp_automatization/tree/main/mavisp_templates/) to capture interactions not yet reflected in PPI databases.

We also use other methods to identify four different classes of short linear motifs (BRCT, LIR, BH3 and UIM) in our target proteins. Depending on the type, we use a combination of simple regular expression matching, a method designed by us for structure-based identification of short linear motifs SLiMfast (available at https://github.com/ELELAB/SLiMfast) together with another method for predicting changes in secondary structure propensity that may be induced by phosphorylation in the core of putative LIR motifs, phosphoriLIR (https://github.com/ELELAB/phospho-iLIR), or DeepLoc 2.0^116^ for predicting the subcellular localization of the protein, especially useful for BRCT motifs.

### Free energy calculations for STABILITY, LOCAL INTERACTION and LONG-RANGE modules

We applied the BuildModel module of FoldX5 suite^117^ averaging over five independent runs for the calculations of changes in free energy of folding upon amino acid substitution with MutateX and the FoldX5 method. We used the cartddg2020 protocol for folding free energy calculations with Rosetta suite and the ref2015 energy function. In this protocol, only one structure is generated at the relax step and then optimized in Cartesian space. Five rounds of Cartesian space optimization provide five pairs of wild-type and mutant structures for each variant. The change in folding free energy is then calculated on the pair characterized by the lower value of free energy for the mutant variant, as described in the original protocol^118^.

We used MutateX to calculate changes in binding free energy for the LOCAL INTERACTION module using the BuildModel and AnalyzeComplex functions of FoldX5 suite and averaging over five runs. With Rosetta, we used the flexddg protocol as implemented in RosettaDDGPrediction and the talaris2014 energy function. We used 35,000 backrub trials and a threshold for the absolute score for minimization convergence of 1 Rosetta Energy Unit (REU). The protocol then generates an ensemble of 35 structures for each mutant variant and calculates the average changes in binding free energy. We used Rosetta 2022.11 version for both stability and binding calculations. In the applications with RosettaDDGPrediction the Rosetta Energy Units (REUs) were converted to kcal/mol with available conversion factors^118^. We also applied RaSP using the same protocol provided in the original publication^56^ and adjusting the code in a workflow according to MAVISp-compatible formats (https://github.com/ELELAB/RaSP_workflow). We have included data on 131 complexes at the date of 16/10/2025 (https://osf.io/y3p2x/).

For the calculations of allosteric free energy, we used the structure-based statistical mechanical model of allostery (SBSMMA)^119,120^ implemented in AlloSigMA2^64^. The model describes the mutated variants as ‘UP’ or ‘DOWN’ mutations depending on difference in steric hindrance upon the substitution. We followed a recently updated and benchmarked protocol^65^. In brief, we classified as uncertain those variants for which the absolute changes in the volume of the side chain upon the amino acid substitution was lower than 5 Å^3^, as recently applied to p53^38^. As a default, we considered as having an effect only variants that were exposed to the solvent (≥25% relative solvent accessibility of the side chain), with associated changes in absolute value of allosteric free energy larger than 2 kcal/mol and considered as remote response sites those that were at a distance higher than 5.5 Å from the mutation site, considering all heavy atoms, and which belongs to pockets as identified by Fpocket^121^ (see workflow at https://github.com/ELELAB/MAVISp_allosigma2_workflow/)

### Efoldmine

The EFOLDMINE module, integrated within the simple mode of MAVISp, predicts residues with early folding propensity using the EfoldMine tool^72^. Trained on residue-level hydrogen/deuterium exchange nuclear magnetic resonance (HDX NMR) folding data from the Start2Fold database^122^, this tool uses secondary structure propensity and backbone/side-chain dynamics in a support-vector machine algorithm to predict early folding regions based on the target’s sequence. In MAVISp, we incorporated EfoldMine to determine whether point mutations in variants fall within the predicted early folding regions, using a threshold of 0.169 to define residues involved in early folding events as suggested by the developers of the method^72^ and considering only regions with a minimum length of three early folding residues to exclude isolated peaks.^71^.

### FUNCTIONAL SITE module

The FUNCTIONAL SITES module aids the identification of variants that might impact cofactor binding sites or active site residues, as well as the residues within the second coordination sphere with respect to active site residues of enzymes or their corresponding binding sites. It is based on a contact analysis performed with the Arpeggio software^123^. Before the analysis, the model structure is subjected to energy minimization with Conjugate Gradients^124^ in 50 steps, using the MMFF94 force field^125^, a van der Waals cutoff of 0.1, an interacting cutoff of 5.0 Å, and a physiological pH of 7.4. Subsequently, the output is further preprocessed to exclude clashes and proximal contacts (https://github.com/ELELAB/mavisp_accessory_tools).

### Molecular dynamics simulations for MAVISp ensemble mode

We used either previously published^37,38,126–130^ or newly collected one microsecond all-atom molecular dynamics simulations performed using the CHARMM22* or CHARMM36m force fields^131^. All the simulations have been carried out in the canonical ensemble after a final equilibration steps and using explicit solvent and periodic boundary conditions. The templates files used for the simulations are provided in OSF (https://osf.io/y3p2x/).

Ensembles generated using simulations are then subject to quality control, either using Mol_Analysis^132^ or MetaD_Analysis (https://github.com/ELELAB/MetaD-Analysis) tools.

As a first example of how we intend to use metadynamics data for the FUNCTIONAL_DYNAMICS module we used the simulations from TP53 where the effects of amino acid substitutions on an interface for protein-protein interaction (residues 207-213) was investigated. We used a collective variable based on distances between two residues (D208-R156) that were effective in capturing open (active) and closed (inactive) conformations of the loop. See repositories associated with the enhanced sampling simulations of TP53^38^. All the newly generated trajectories will be deposited as different entries in OSF, and the link is reported in the metadata on the MAVISp webserver. At the date of 01/11/2024, we have included 45 protein targets in the *ensemble mode* using as source of ensemble mostly unbiased MD simulations of 500 ns or one-μs, as detailed in the corresponding metadata on the MAVISp webserver. In some cases, we included ensembles generated by a coarse-grain model of flexibility or using the conformation provided by NMR structures from the PDB (see INPUT STRUCTURES tables in https://osf.io/y3p2x/).

### Protein Structure Networks and path analysis for MAVISp ensemble mode

In the ensemble mode we apply a module building upon the simple mode LONG_RANGE module. It uses AlloSigma2-PSN (https://github.com/ELELAB/MAVISp_allosigma2_workflow/) where we constructed an atomic-contact PSN on the full trajectories using PyInteraph2^66^. Pairs of residues were retained only if their sequence distance exceeded Proxcut threshold of 1 and their edge calculations remained within less than 4.5Å, based on the thresholds described in PyInteraph2^66^. We retained edges with an occurrence greater than Pcrit threshold of 50% across the ensemble frames, weighted on the interaction strength Imin of 3.

Subsequently, we used the path_analysis function of PyInteraph2 to identify the shortest paths of communication between each pair of AlloSigMA2^64^ predicted mutation and respective response sites, using a minimum distance threshold of 5.5 Å and retained paths that were four residues or longer.

### CABS-flex ensembles for MAVISp ensemble mode

We used the coarse-grained CABS-flex 2.0 method and software^50^ as a part of a Snakemake^133^ pipeline, available at https://github.com/ELELAB/MAVISp_CABSflex_pipeline. The pipeline includes the possibility to tune the calculations by different restraints, secondary structure definition, ligand binding and more. It also contains a quality control step to evaluate the secondary structure content of the generated structures with respect to the starting one, using DSSP^134^ and the SOV-refine score^135^.

### Variant Effect Prediction

We used DeMaSk^74^, GEMME^14^, EVE^75^, REVEL^73^ and AlphaMissense^26^ as predictors for the effect of any possible amino acid substitution to natural amino acids, on the full protein sequence of the main UniProt^136^ isoform of each protein. We used available default parameters for each method unless noted otherwise. We used the standalone version of DeMaSk as available on its public GitHub (commit ID 10fa198), with BLAST+ 2.13.0. We followed the protocol available on GitHub: we first generated the aligned homologs sequence file by using the demask.homologs module and then calculated fitness impact predictions. Finally, we classified as loss-of-fitness those variants having a DeMaSk delta fitness score in absolute value lower or equal to - 0.25, gain-of-fitness if the score is higher than 0.25, and neutral otherwise (**Supplementary Text S3**). We used the available online webserver to obtain variant effect predictions with GEMME, upon setting the number of JET iterations to 5, to obtain more precise results. ^126^ We have classified variants having a GEMME score <= -3 as damaging, and neutral otherwise. Thresholds were selected according to our benchmarking (**Supplementary Text S3).** To obtain EVE scores, we have used the scripts, protocol and parameters available on the EVE GitHub (commit iD 740b0a7) as part of a custom-built Snakemake^133^–based pipeline, available at https://github.com/ELELAB/MAVISp_EVE_pipeline Using EVE first requires building a protein-specific Bayesian variational autoencoder model, which learns evolutionary constraints between residues from a multiple sequence alignment. In the current MAVISp release, we generated such alignments using EVcouplings^137^, using the Uniref100^138^ sequence database released on 01/03/2023, by keeping sequences with at least 50% of coverage with the target protein sequence, alignment positions with a minimum of 70% residue occupancy, and using a bit score threshold for inclusion of 0.5 bits with no further hyperparameter exploration. We then used our pipeline to perform model training, calculation of the evolutionary index, and used a global-local mixture of Gaussian Mixture Models to obtain a pathogenicity score and classification. We have used pre-computed REVEL scores for variants as available in dbSNFP^139,140^, accessed through myvariants.info^141,142^, as implemented in Cancermuts. We have classified as damaging variants that have a REVEL score larger or equal to 0.5^143^. We included AlphaMissense pathogenicity prediction scores and classification as available by the dataset of prediction for all possible amino acid substitutions in UniProt canonical isoforms, release version 2^144^.

### Annotations from experimental data for EXPERIMENTAL_DATA module

We developed Python scripts to identify the overlap in coverage between the Mave database (MaveDB)^92^ and MAVISp, and to retrieve the score sets associated with the shared entries from the MaveDB^92^ database through their API (https://api.mavedb.org/docs). Where available, we also extracted information on methods and classification thresholds. For entries where this information was incomplete, the corresponding publications were manually reviewed to extract thresholds for variant classification.

The ProteinGym^91^ repository was locally downloaded from GitHub, and a custom Python script was used to process the datasets based on the reference files provided in the repository. The datasets used for the analysis contained the experimental scores and the classification provided by the authors either based on the median of the score distributions or via manual annotation. The scores and their classifications were then integrated into the final database file generated by MAVISp. The aggregated scores, along with their classifications, were compiled into the final database file produced by MAVISp through a module dedicated to the experimental data.

### Identification of RefSeq identifiers

To ensure the correct RefSeq annotations in MAVISp, we implemented a Python tool, *compare_seq.py* (https://github.com/ELELAB/mavisp_accessory_tools/), to verify the sequence identity between the canonical UniProt sequence used in our analyses and the corresponding RefSeq protein identifier to be used for the ClinVar search. The Uniprot sequences were retrieved using the UniProt REST API, while the RefSeq protein sequences were fetched from the NCBI Entrez Protein database. We implemented a global pairwise alignment using the Biophyton pairwise2 module with the globalxx scheme to assess sequence identity. Each comparison was classified as either an exact match, a mismatch (identity <100%), or unresolved due to missing or unresolvable sequences. To improve performances, the analyses were parallelized using multi-threading via Python concurrent.futures. The results were logged into structured CSV reports for consultation. This allows data managers to identify exisiting entries in MAVISp with RefSeq identifiers inconsistent with provided UniProt accession code and assign them to biocurators for entry review.

Additionally, we provide the biocurators with a Python-based script (*uniprot2refseq,* l https://github.com/ELELAB/mavisp_accessory_tools/) that identifies RefSeq IDs for the UniProt canonical protein isoform. For each UniProt AC, we queried the UniProt REST API to obtain RefSeq protein cross-references (NP_* IDs) from the canonical entry in JSON format. Only protein-level RefSeq entries were considered. The canonical UniProt protein sequence was downloaded in FASTA format, and each RefSeq sequence was retrieved from the NCBI Protein database using Biopython and the Entrez API. Pairwise global alignments were performed using the Biopython pairwise2 module and we estimate the percentage sequence identity as the number of identical residues over the length of the longer sequence. Results were saved in tabular format, including UniProt AC, RefSeq ID, and sequence identity. This approach aids the biocurators to identify the RefSeq IDs for the canonical isoform of the protein undebefore starting with the data collection. The script is expected to be used by the biocurators before each run with the MAVISp automatization workflow described below.

### Workflows for automatization and data collection within MAVISp

We provide and maintain two Snakemake workflows for the data collection of the default modules of MAVISp. The first is a Snakemake pipeline to automate MutateX runs as much as possible. It is designed to automatically download the chosen structure(s) from the AlphaFold structural database, or a custom structure input file, when necessary, trim them as requested, and generate desired MutateX folding free energy scans with a predictable directory structure. It only requires as input a csv file with metadata on the desired scan and a configuration file with details on the run to be performed. It is available at https://github.com/ELELAB/mutatex_pipelines/tree/main/custom_collect_scan.

Once such a scan is available, it is possible to use a second Snakemake pipeline, called MAVISp_automatization, which performs most of the steps that are necessary to annotate a protein for a MAVISp simple mode entry. Similarly to the previous pipeline, it only requires metadata on the target protein to be analyzed, as well as a MutateX mutational scan. It generates a dataset that can then be imported into the MAVISp database, except for predictions performed using Rosetta-based methods, since these are much more computationally expensive and need to be performed separately using the RosettaDDGPrediction pipeline^55^. Using a Snakemake pipeline allows to improve efficiency and scalability, allowing to use multi-core system to process several proteins or perform different analyses in parallel. It is available at https://github.com/ELELAB/MAVISp_automatization.

## Data Availability

The data can either be consulted through our web server (https://services.healthtech.dtu.dk/services/MAVISp-1.0/) or as individual CSV files in the OSF repository https://osf.io/ufpzm/. Other raw data and utilities can be found at the MAVISp extended data OSF repository (https://osf.io/y3p2x/) Reports for several proteins are available at https://elelab.gitbook.io/mavisp/.

## Author Contributions (CRediT Classification)

*Conceputalization*: EP, *Data Curation*: MA, MU, KD, LBa, SS, KK, PSB, KM, LBb, TD, AM, LF, EK, AO, AHE, JB, JS, FM, BAF, GT, PB, JV, ML, MT, EP *Formal Analysis*: MA, LBa, MU, TD, ML, EP. *Funding Acquistion:* EP, MA, CC, SI, MN, AR. *Methodology*: MA, MU, LBa, KD, SS, KK, KM, LBb, EK, JB, AP, JV, ML, MT, EP. *Project administration:* EP. *Resources*: PWD, EP, SI, CC. *Software*: MA, KD, KK, SS, LBb, AP, ML, MT, PWS. *Supervision*: EP, ML, MT, SS, MU, MA, LBa. *Validation*: All the coauthors. *Visualization*: MA, MU, KD, LBa, MU, MT. *Writing – Original Draft:* EP, MT. *Writing – Review and Editing*: All the coauthors.

## Acknowledgements

Our research has been supported by Carlsberg Foundation Distinguished Fellowship (CF18-0314), Danmarks Grundforskningsfond (DNRF125), Hartmanns Fond (R241-A33877), LEO Foundation (LF17006), NovoNordisk Fonden Bioscience and Basic Biomedicine (NNF20OC0065262) to E.P. group. Part of the calculations have been supported by a EuroHPC Benchmark Access Grant (EHPC-BEN-2023B02-010) and a EuroHPC Regular Grant (EHPC-REG-2023R01-051) on Discoverer. The work is also supported by A PhD Fellowship from the Danish Data Science Academy (DDSA) to M.A. SI and JS would like to thank the Merkin Institute of Transformative Technologies in Healthcare. AR is supported by The Assar Gabrielsson’s foundation and The Healthcare Board, Region Västra Götaland. PSB is supported by a KBVU Pre-Graduate Fellowship (R361-A21156).

This project is also in part based upon work from COST Action ML4NGP, CA21160, supported by COST (European Cooperation in Science and Technology) which supported a short-term visit for GT in EP group.

KD, MT and EP would like to thank Kresten Lindorff-Larsen, Lasse M. Blaabjerg and Matteo Cagiada for their inputs and feedback to port the RaSP code within our workflow. We also would like to thank the group of Stefano Vanni for providing published MD trajectories for analysis within the ensemble mode of MAVISp. We would like to thank Jeppe Samuelsen, Kledi Salla, Amalie Drud Nielsen, Julie Bruun Brockhoff, Laura Mattioli, Zeming Hou, Subayan Akhuli and Ona Saulianskaite, Subhayan Akhuli, Martina Bellini, Eirini Giannakopoulou, Beatrice Drago, Eszter Toldi, Laura Kappel, Alessia Campo, Angeliki Vliora, Vinit Nilesh Vasa, Edene Levine, Konstantina Gkopi, Alicia Llorente for preliminary work on targets reported in the database.

