## Supplementary Text S1 for "MAVISp: A Modular Structure-Based Framework for Protein Variant Effects"

**Comparison of RaSP and Rosetta in the consensus approach used for the STABILITY module of MAVIsp**

We have benchmarked the RaSP method^1^ for its suitability to be used as an alternative to the Rosetta *cartΔΔG2020* protocol^2^ in the context of the high-throughput data collection for MAVISp *simple mode*. RaSP is a machine-learning method to predict change of folding free energy upon mutation that has been trained against predictions performed with Rosetta. The main advantage of applying RaSP over Rosetta in the context of MAVISp, considering the large size of our dataset, is its much lower computational cost; furthermore, it has been demonstrated to perform accurate predictions on AlphaFold structures as used in MAVISp for stability calculations. According to the original RaSP publication^1^, the method displays a Pearson correlation coefficient of 0.82 and mean absolute error (MAE) of 0.73 kcal/mol with respect to much more computationally expensive Rosetta calculations on a test dataset of 10 proteins that were not part of the training set. As our dataset is significantly larger, we performed a similar benchmarking by considering results from Rosetta calculations as our reference and by checking its effect on the MAVISp stability classification. Code and results for this benchmarking are available at <https://github.com/ELELAB/MAVISp_RaSP_benchmark>

For our benchmarking, we have used values of free energy changes for RaSP and Rosetta available in MAVISp. The former were obtained by using our RaSP framework available at <https://github.com/ELELAB/RaSP_workflow>, which is derived from original RaSP code available at [https://github.com/KULL-Centre/_2022_ML-ΔΔG-Blaabjerg/](https://github.com/KULL-Centre/_2022_ML-ddG-Blaabjerg/). The latter were calculated using our RosettaDDGPrediction pipeline^3^ and Rosetta nightly build v. 2022.11. Our dataset consists of proteins included in the MAVISp database as of 20/12/2023. In order to avoid circular bias, we made sure to remove any proteins that were used in the training set of RaSP. We obtained the training dataset for RaSP from the RaSP GitHub repository. As this was a list of Protein Data Bank (PDB^4^) identifiers, we retrieved the corresponding UniProt accession codes for each protein in the structures using the PDB API. This final list of proteins was used to filter the MAVISp data, which left us with 181 proteins out of the 203 proteins in the MAVISp database. After filtering out substitutions where we did not have data for both RaSP and Rosetta, we were left with 70,446 observations (amino acid substitutions) in 179 different proteins. It should be noted that the dataset under consideration is still somewhat uneven, as the 23 proteins with the largest number of mutations (12% of the dataset) still constitute 50% of the total number of observations. Furthermore, in this dataset, the observations are not equally distributed among amino acid types, either considering mutations to or from a certain type (Figure S1.1). For instance, tryptophan is notably underrepresented in both cases. While this might bias the following analysis to some extent, it is representative of the MAVISp dataset which is the focus of this investigation.

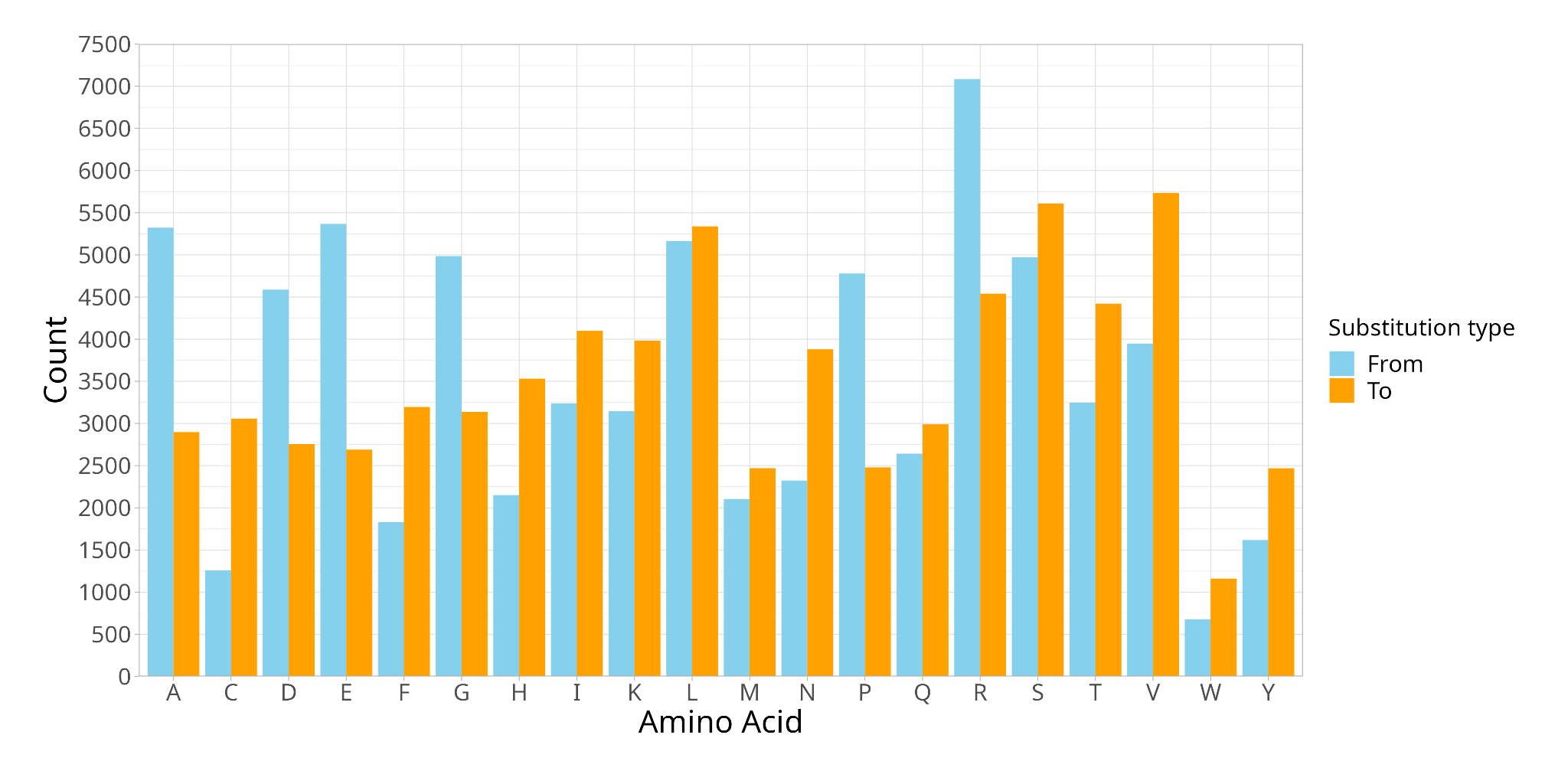

***Figure S1.1*** *Distribution of the different types of amino acid substitutions. From is the type of the wild-type amino acid. To is the type of the mutated amino acid.*

To evaluate the performance of RaSP, mirroring the original publication, we calculated Pearson correlation coefficient and MAE between the Rosetta and RaSP datasets, obtaining 0.73 and 0.37 kcal/mol, respectively (Figure S1.2). We obtained a slightly lower correlation and significantly lower mean absolute error with respect to the values reported in the original publication of RaSP. We noticed a number of outliers in our dataset that feature extremely high ΔΔG values for Rosetta and much more modest values for RaSP; RaSP values top at 11.9 kcal/mol, while the Rosetta dataset features a number of data points well above 15 kcal/mol (up to 62.3 kcal/mol) (Figure S1.3). To assess the effect of such outliers on the final performance measures, we also calculated the Pearson correlation coefficient and MAE without them. We removed the outliers from the dataset by filtering out data points with a Rosetta value >= 15 kcal/mol. This removed 143 out of the 70446 observations (0.2%).We obtained a Pearson correlation of 0.74 and MAE of 0.35 kcal/mol. Since these outliers are a small fraction of the dataset, the Pearson correlation and MAE were largely unaffected.

To check whether the 23 proteins with the greatest number of mutations affect performance, the Pearson correlation coefficient and MAE were also calculated without them. Here we obtained a Pearson correlation coefficient of 0.73 and MAE of 0.4 kcal/mol, showing that they do not skew the general performance.

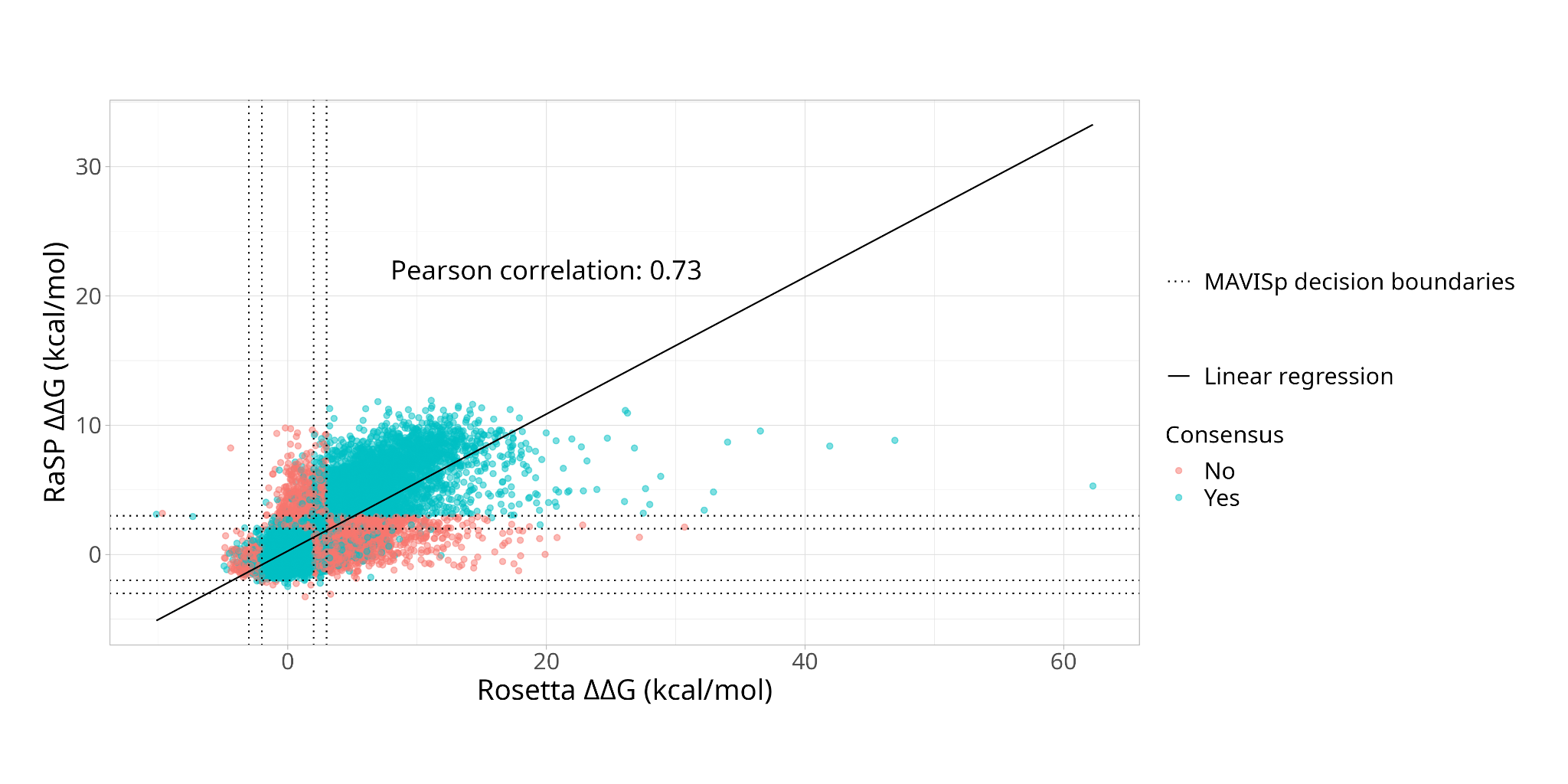

***Figure S1.2*** *Scatter plot of the free energy changes in kcal/mol obtained using RaSP and Rosetta respectively. Each point corresponds to an amino acid substitution. Blue dots are the cases where the MAVISp classification is the same for both of them, and red dots are the cases where they are not classified as the same. For clarification, the decision boundaries of MAVISp have been drawn as dotted lines.*

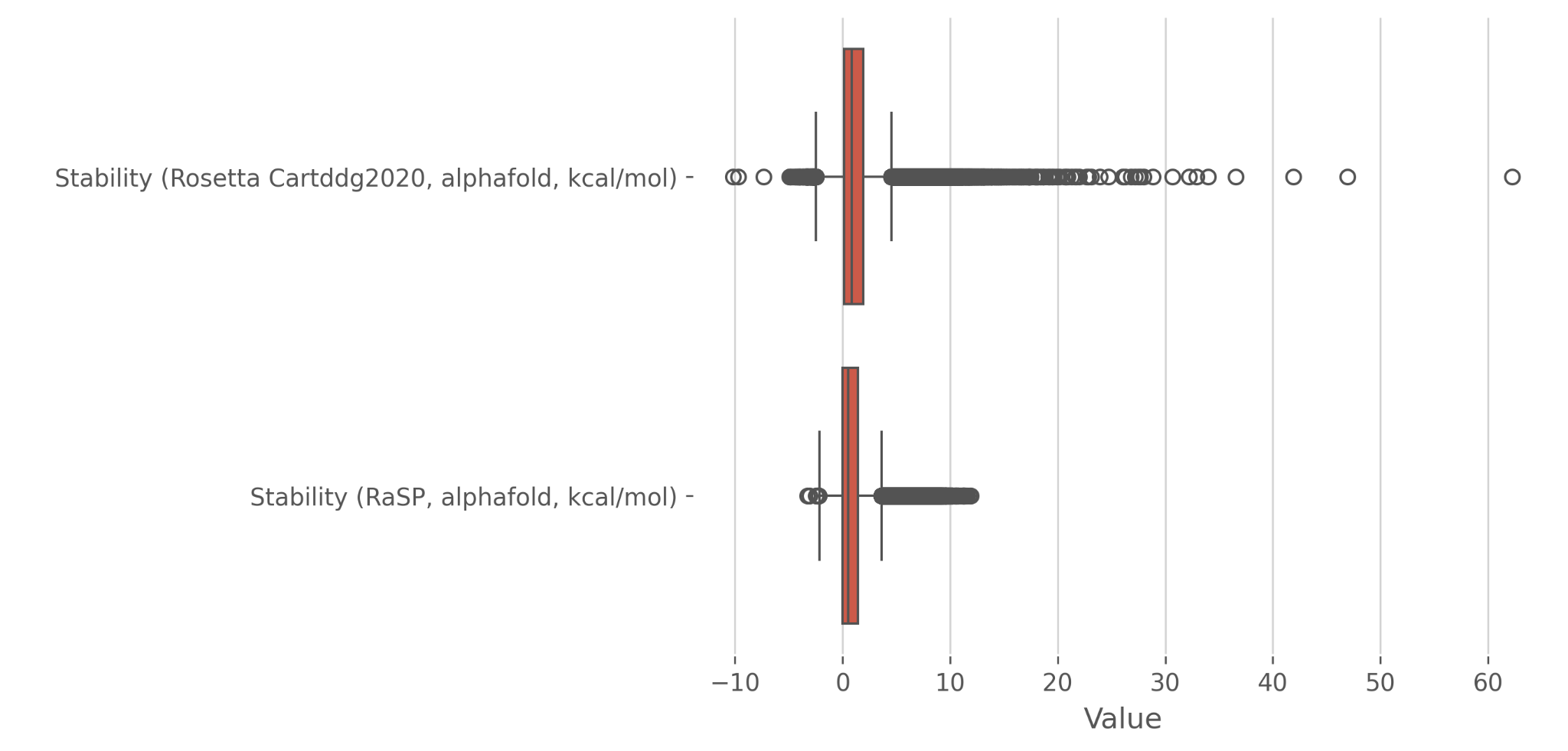

***Figure S1.3*** *Boxplot of the distribution of Rosetta and RaSP predictions illustrating both the high values predicted by Rosetta and the overall lower scores assigned by RaSP.*

Next, we used both RaSP and Rosetta, each together with FoldX^5,6^ predictions of changes of folding free energy upon mutation, to classify the effect of mutations according to the MAVISp stability module. This uses a consensus approach between RaSP or Rosetta and FoldX results, as also explained in the main text. Briefly, for each method, we consider the following classification for any given mutation:

- Destabilizing if ΔΔG ≥ 3 kcal/mol
- Stabilizing if ΔΔG ≤ -3 kcal/mol
- Neutral if -2 < ΔΔG < 2 kcal/mol
- Uncertain otherwise (-2 ≤ ΔΔG < -3 | 2 ≤ ΔΔG < 3 kcal/mol)

Finally, if the two methods return the same classification we assign that classification to the mutation; conversely, if they disagree, the mutation is classified as Uncertain. In the following text, we will refer to the “RaSP protocol” or “Rosetta protocol” as the outcome of this process, either using RaSP or Rosetta, respectively, together with FoldX, as detailed.

Both the RaSP and Rosetta protocols classify the majority of mutations as Neutral (Figure S4). Our Rosetta protocol only predicts 6 mutations as stabilizing, with 0 for the RaSP protocol (Figure S1.4 and S1.5. Our analysis (Figure S1.5) shows that the majority of mutations are classified in the same way for both the Rosetta and RaSP protocols. This is mostly due to a large overlap between the Neutral and Uncertain classes. Notably, 4367 mutations are classified as Destabilizing for both protocols, nonetheless 2417 are classified as Destabilizing for the Rosetta protocol and uncertain for the RaSP protocol. This is only partially offset by 621 mutations, which are Destabilizing for the RaSP protocol while are classified as Uncertain by Rosetta.

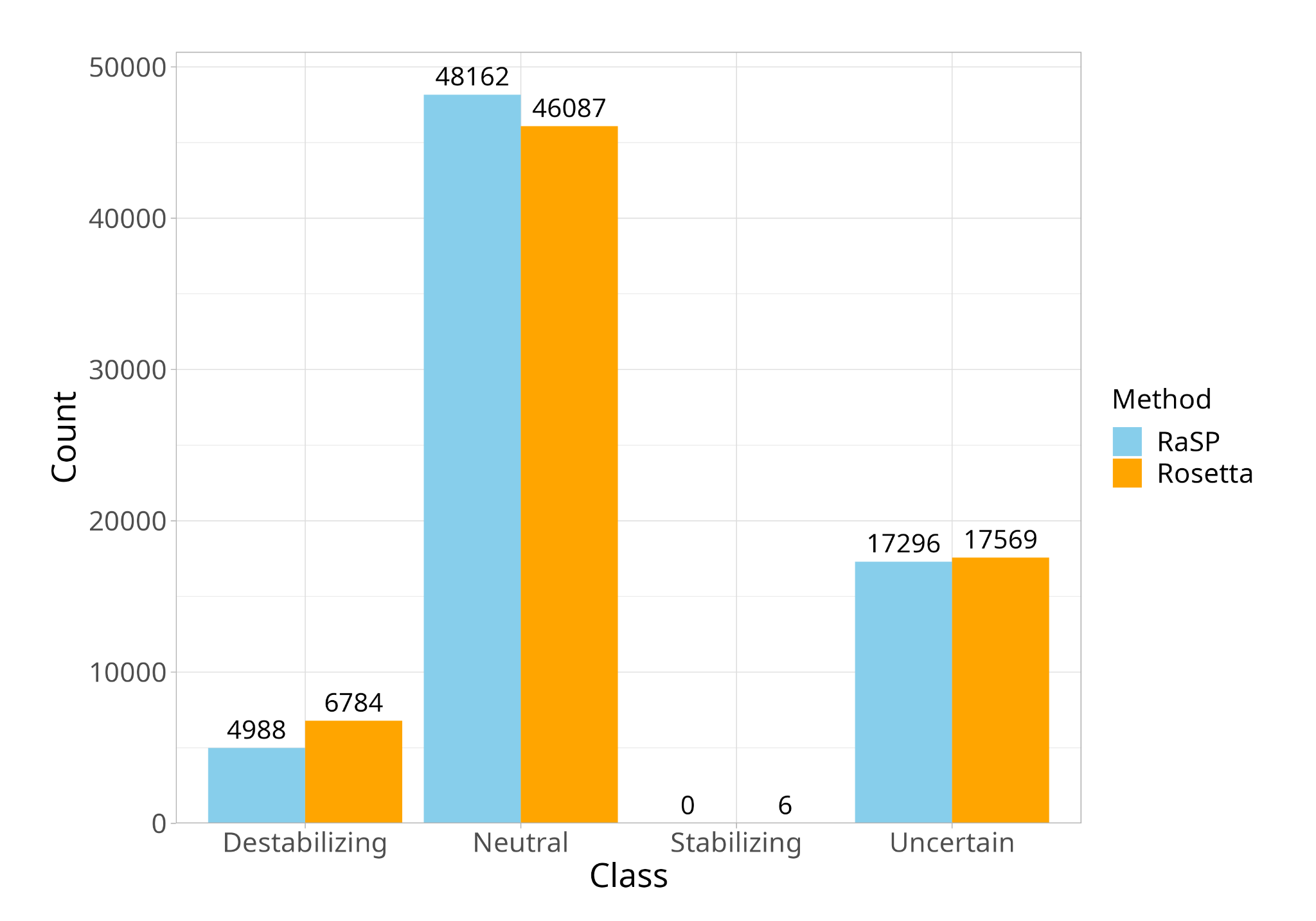

***Figure S1.4*** *Number of amino acid substitutions in each class for RaSP vs. Rosetta protocol.*

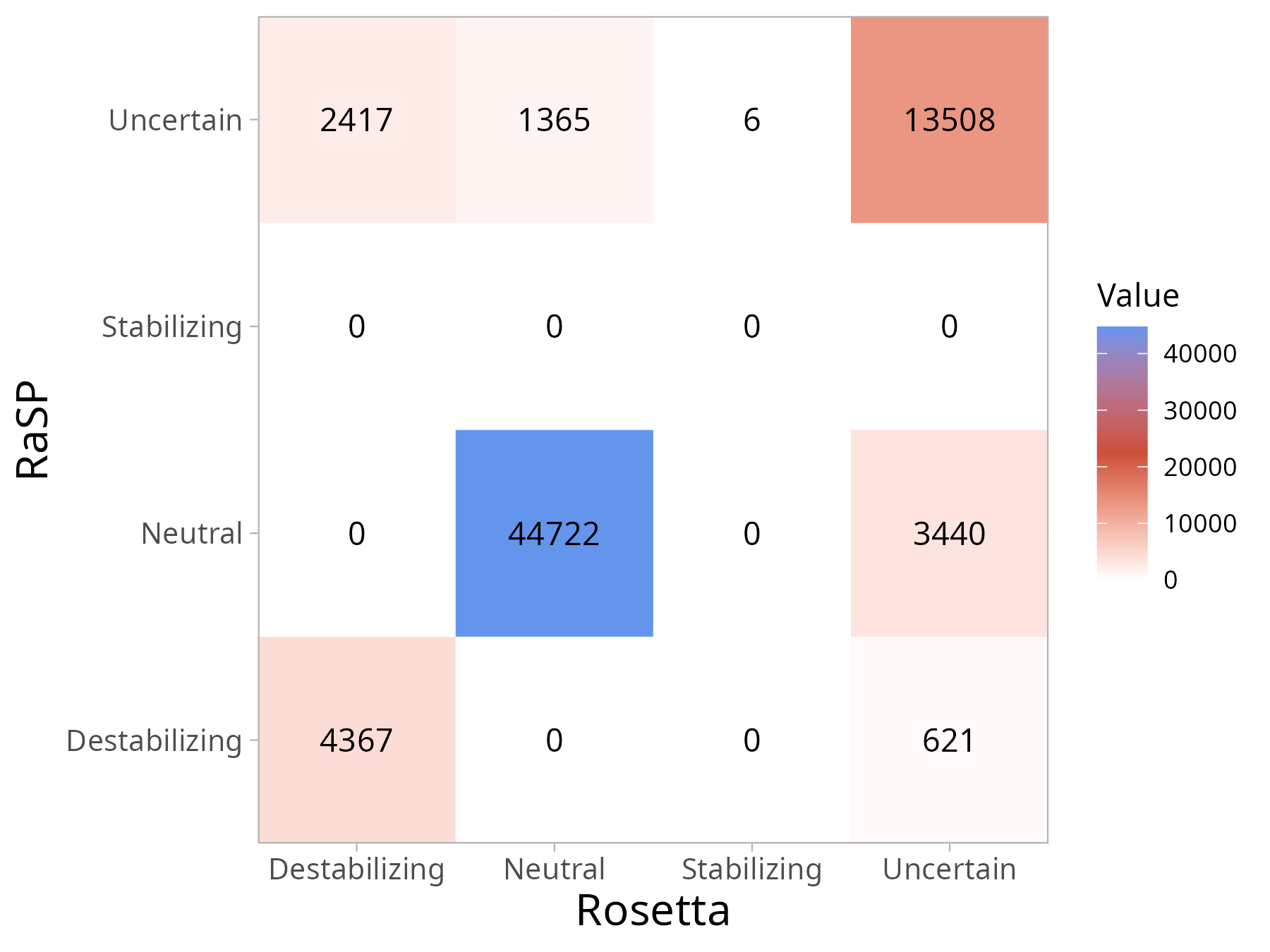

***Figure S1.5*** *Confusion matrix between the RaSP and Rosetta protocol.*

For the purposes of MAVISp, it is important to not misclassify the substitutions to a degree that will compromise the quality of the prediction. The disagreement between the two classifications mainly concerns uncertain classifications. For the most part, the RaSP protocol classifies substitutions as uncertain, which are otherwise classified by Rosetta. In the case that the RaSP protocol classifies an uncertain substitution as something else, it is mostly as neutral, and more rarely as destabilizing or stabilizing. Hence, the reliability of the conclusive classifications (Neutral, Destabilizing, Stabilizing) are not largely compromised by using the RaSP protocol to classify the substitutions. However, the classification that has the most impactful biological implications in the MAVISp framework is the one of Destabilizing mutations; in this case, the RaSP protocol classifies a number of mutations differently with respect to the Rosetta protocol.

We also verified if there exists some bias in the classification between the Rosetta and RaSP protocol for certain types of mutations, i.e. if the RaSP protocol systematically misclassifies certain types of substitutions. We looked at different aspects of the substitutions such as the amino acid types and the solvent accessibility of the site. In each of the cases, we defined accuracy as the number of correct classifications in which the Rosetta protocol classification was the same as the RaSP protocol, divided by the total number of observations in the subset.

First, we looked for bias in the classification considering amino acid types. Each amino acid substitution would be defined as both a “from” substitution and a “to” substitution. As an example, the mutation A54W would be categorized as “from A” and “to W”. We calculated accuracy for each “from” and “to” residue types. Data for this analysis was available for all the 70446 observations.

First, looking broadly at the accuracy of the amino acid substitutions, both to and from each amino acid, no single amino acid type sticks out (Figure S1.6). This indicates that no particular wild-type or mutated residue types are significantly more likely than others to cause misclassification.

The original RaSP publication identified larger errors for glycine substitutions and when changing to proline, when comparing results for Rosetta and RaSP. Using the consensus approach, we do not see a significant bias towards glycine or proline substitutions or other amino acids for that matter.

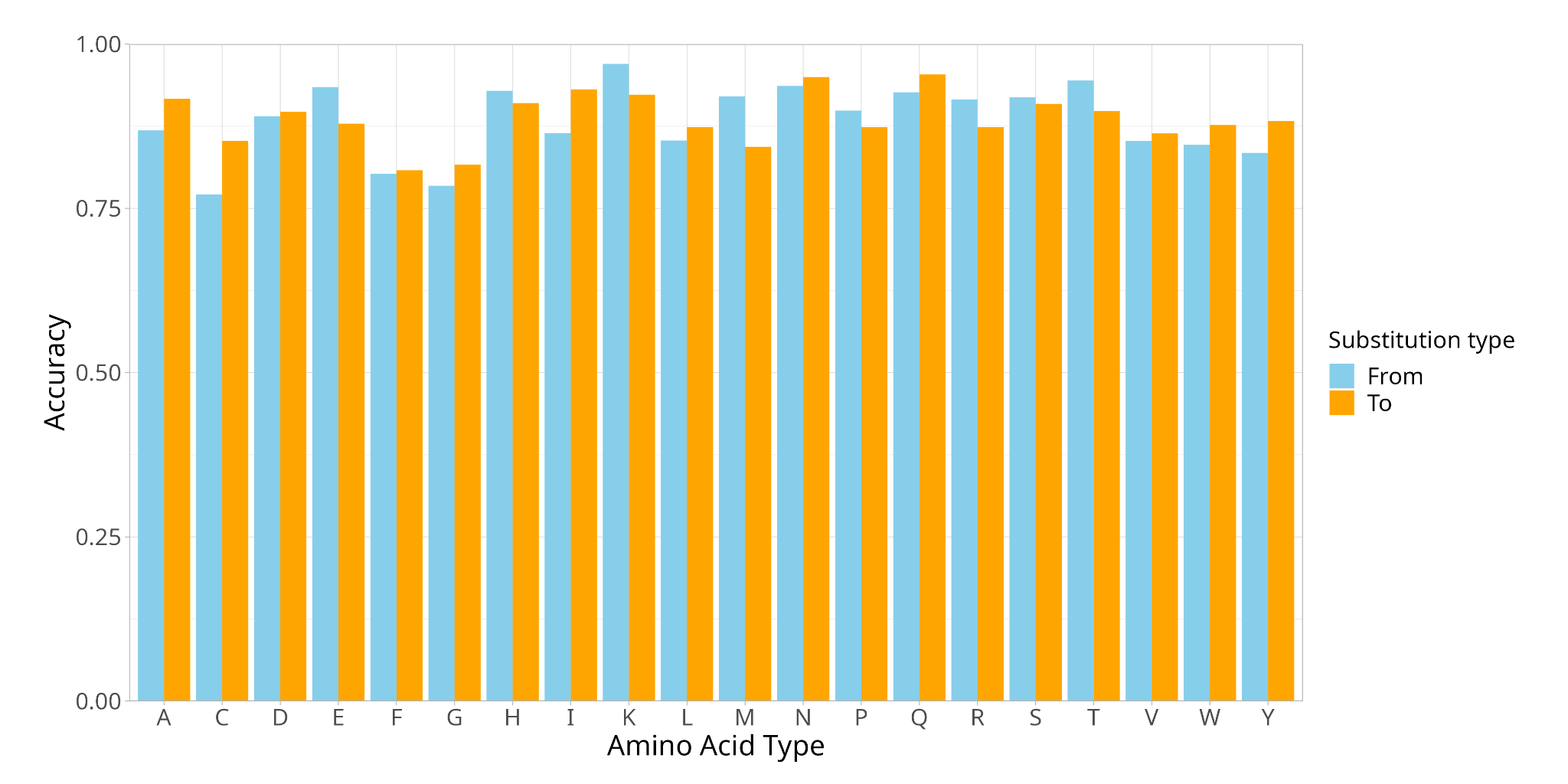

***Figure S1.6*** *Accuracy of amino acid substitutions. Amino acid substitutions are categorized into From and To. From is the wild-type amino acid type, and To is the mutated amino acid type.*

The original RaSP publication also identifies better accuracy for solvent exposed sites with respect to buried sites. To investigate this, we used the per-residue, side-chain relative solvent accessible surface area (SASA) values available in MAVISp. These were calculated on the wild-type protein structures collected in MAVISp, using NACCESS^7^. The observations were divided in two categories based on whether the Rosetta and RaSP protocol gave the same classification. A kernel density estimator was fitted to these two groups, with the assumption that the curves would be very similar if there was no bias. We found that the group of mutations in which the two classifications did not agree is made of a larger number of buried sites (low SASA), while the group in which the two classifications agree is enriched in more solvent-exposed residues (Figure S1.7).

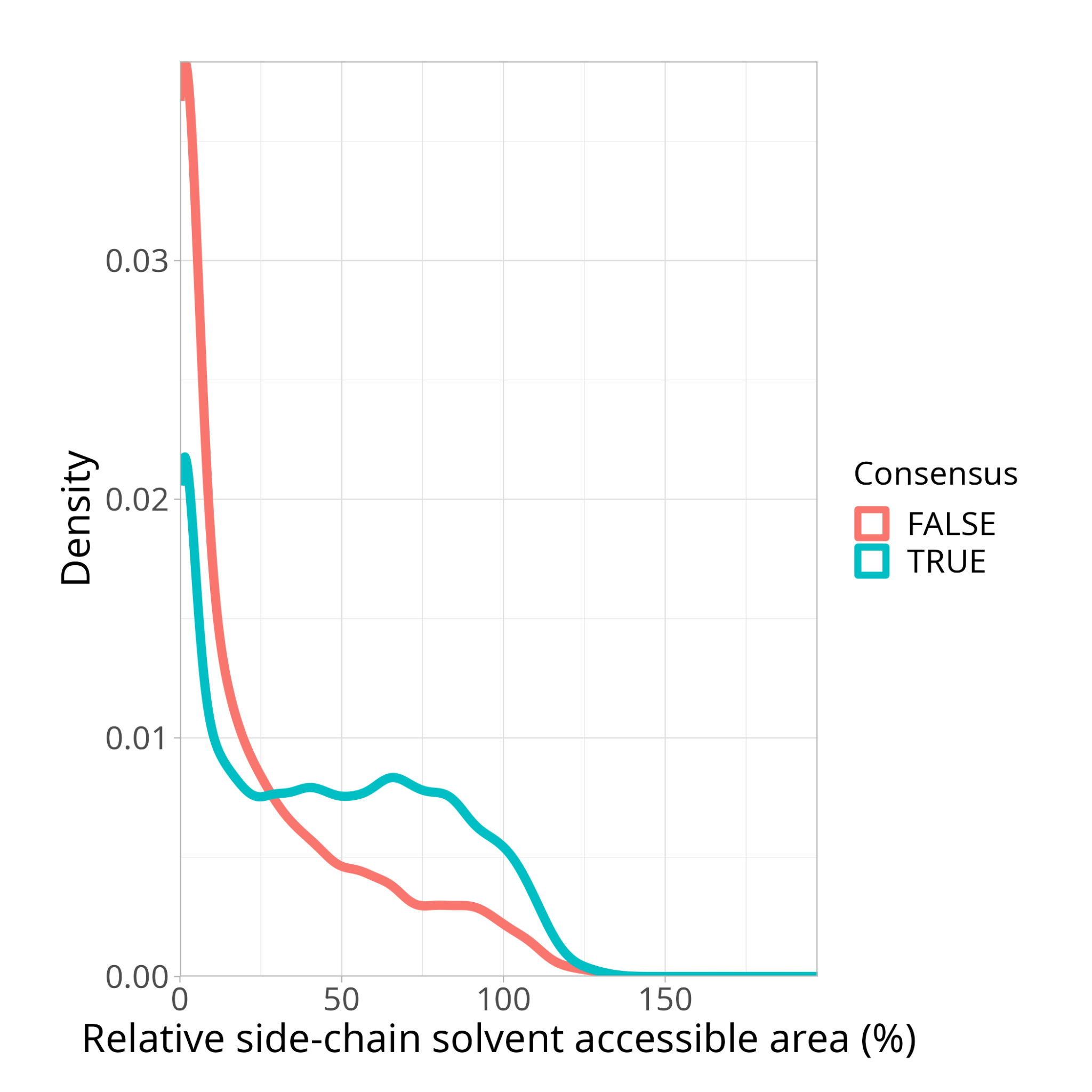

***Figure S1.7*** *Density plot SASA.*
