## Supplementary Text S2 for "MAVISp: A Modular Structure-Based Framework for Protein Variant Effects"

**Text S2 - Detailed description of the methodology used by the PTM module**

The PTM module is designed to classify the effect on mutations on sites subjected to post-translational modifications (PTMs). As different PTM types are very diverse in terms of chemical markup and effects, our long-term plan is to design independent sub-modules for each one independently.

For the first release, we only consider the most well-studied and relevant type of PTM - that is phosphorylation. Phosphorylation can affect several aspects of a protein and biological pathways. Here we focus in particular on three aspects, and classify the effect of each mutation on phosphorylation sites according to its ability to affect i) protein regulation by phosphorylation itself ii) protein structural stability stability iii) function, in terms of binding with partners. This means that the module provides three independent classifications per mutation site.

The known phosphorylation sites are retrieved using the Cancermuts Python package [1], which uses information from a local copy of the PhosphoSitePlus database [2] which is updated yearly. Once we have annotated the experimentally known PTM sites, we perform classification for the three different aspects separately. Each of them uses further data and a custom workflow to derive an annotation, as explained in the sections below.

*Annotation on regulation*

This part annotates the mutations on their potential effect to disrupt the regulation of protein activity or function when a PTM site is mutated. It uses the following information:

- primary sequence of wild-type and mutant variants
- relative side-chain solvent-accessible surface area, as calculated on a structural model of the wildtype protein using NACCESS [3] in simple mode. In ensemble mode, the average of the same value over the ensemble is considered instead.

It follows the logic detailed in Figure S3.1. First, it considers whether any mutated position is a PTM site. If it is, but the mutation could lead to retaining the phosphorylation site (S to T or T to S), it is annotated as Neutral. If this is not the case, and the mutation site is solvent-exposed (SAS>= 25%) and the mutation is not to/from one of the phosphorylatable residues (T, S, Y), it is annotated as Damaging. Otherwise, it is annotated as Uncertain.

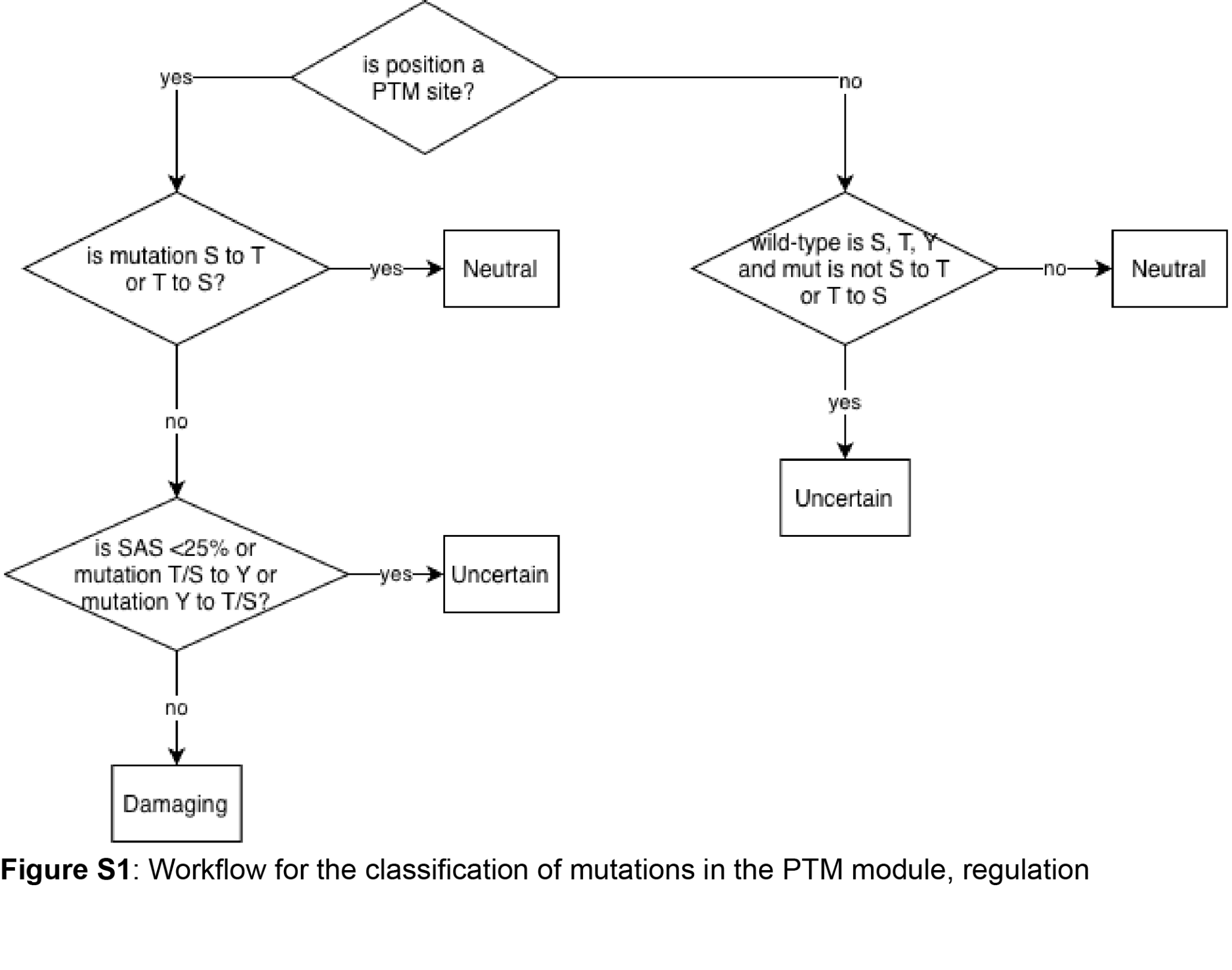

**Figure S3.1**. Workflow for the annotations of variant effects in the PTM REGULATION module

*Annotations on stability*

This part annotates the mutations on their potential effect to alter the stability of the protein in the same or different way as phosphorylation would, when a phosphorylation site is mutated. In short, it provides a prediction of the mutation ability to retain or not the effect that phosphorylation would have on regulating protein stability.

The PTM STABILITY module uses the following information:

- primary sequence of wild-type and mutant variants
- change of folding free energy upon mutation (DDGmut) predicted by FoldX [4], with its consequence classification according to thresholds defined for the STABILITY module
- change of folding free energy upon phosphorylation of wild-type residue (DDGphospho), with its consequence classification according to thresholds defined for the STABILITY module

The module tries to assess whether the phosphorylation and mutation have the same effect on stability (as in stabilizing, neutral, or destabilizing the protein structure); if the effect is different, the mutation is considered damaging.

More in detail, the module follows the workflow outlined in figure S3.2. First, it checks if the mutation site has a potentially phosphorylatable residue type; if it’s not, the mutation is classified as Neutral. Furthermore, if the PTM site is not a known phosphorylation site or if the changes in free energy values are not available (for instance, if the site is in a non-structured region), the mutation site is not classified. If the stability predictions are available, but either DDGmut or DDGphospho are classified as Uncertain, then the final classification is also Uncertain. Finally, if DDGmut and DDGphospho have the same classification, the mutation is classified as Neutral; Otherwise, it is classified as damaging if they differ.

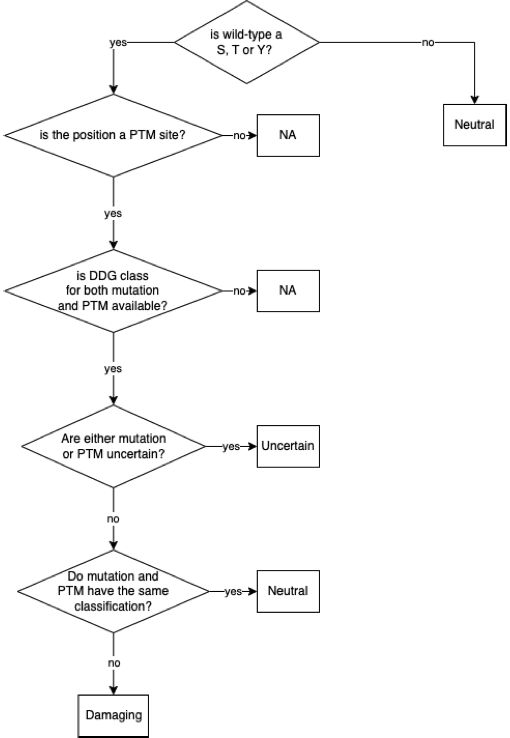

**Figure S3.2**. Workflow for the classification of mutations in the PTM STABILITY module

*Annotations on function*

This part annotates the mutations on their potential effect on altering protein function when a PTM site is mutated. In particular, it focuses on the fact that a mutation that abolishes a phosphorylation site might also affect binding with protein partners when the phosphorylation is essential for binding. It focuses on cases where phosphorylation occurs in a known short linear motif. It uses the following information:

- primary sequence of wild-type and mutant variants
- change of binding free energy upon mutation (DDGmut) predicted using FoldX [4], with its consequence classification according to thresholds defined for the local_interactions module
- change of binding free energy upon phosphorylation of wild-type residue (DDGphospho), with its consequence classification according to thresholds defined for the local_interactions module
- relative side-chain solvent-accessible surface area, as calculated on a structural model of the wildtype protein using NACCESS [3] in simple mode. In ensemble mode, the average of the same value over the ensemble is considered instead
- whether the phosphorylation site is part of a potential short linear motif: this is provided by Cancermuts [1] using ELM [5]
- list of phospho-motifs: this is a hand-made curation of known ELM [5] linear motifs for which binding to a partner is dependent on the phosphorylation state of the motif. It is available on the MAVISp GitHub.

The module follows a workflow described in Figure S3.3, and follows a design similar to that of the stability classification. First, it checks if the mutation site has a potentially phosphorylatable residue type; if not, the mutation is classified as Neutral. Further, if the PTM site is not a known phosphorylation site, the mutation site is not classified. If either DDGmut or DDGphospho are not available, MAVISp checks whether i) the site is part of a known phospho-SLIM considering our custom-made curation and ii) the wild-type residue is solvent accessible (SAS >= 25%). If both conditions are satisfied, the mutation is classified as Potentially Damaging, and Uncertain otherwise.

If both DDGmut and DDGphospho are available, they are used to perform the classification instead. If they have the same classification, then the mutation is considered Neutral, and Damaging otherwise. It should be noted that the classification of changes in free energy doesn’t have an Uncertain case, and therefore there’s no need to handle it as it is the case in stability classification.

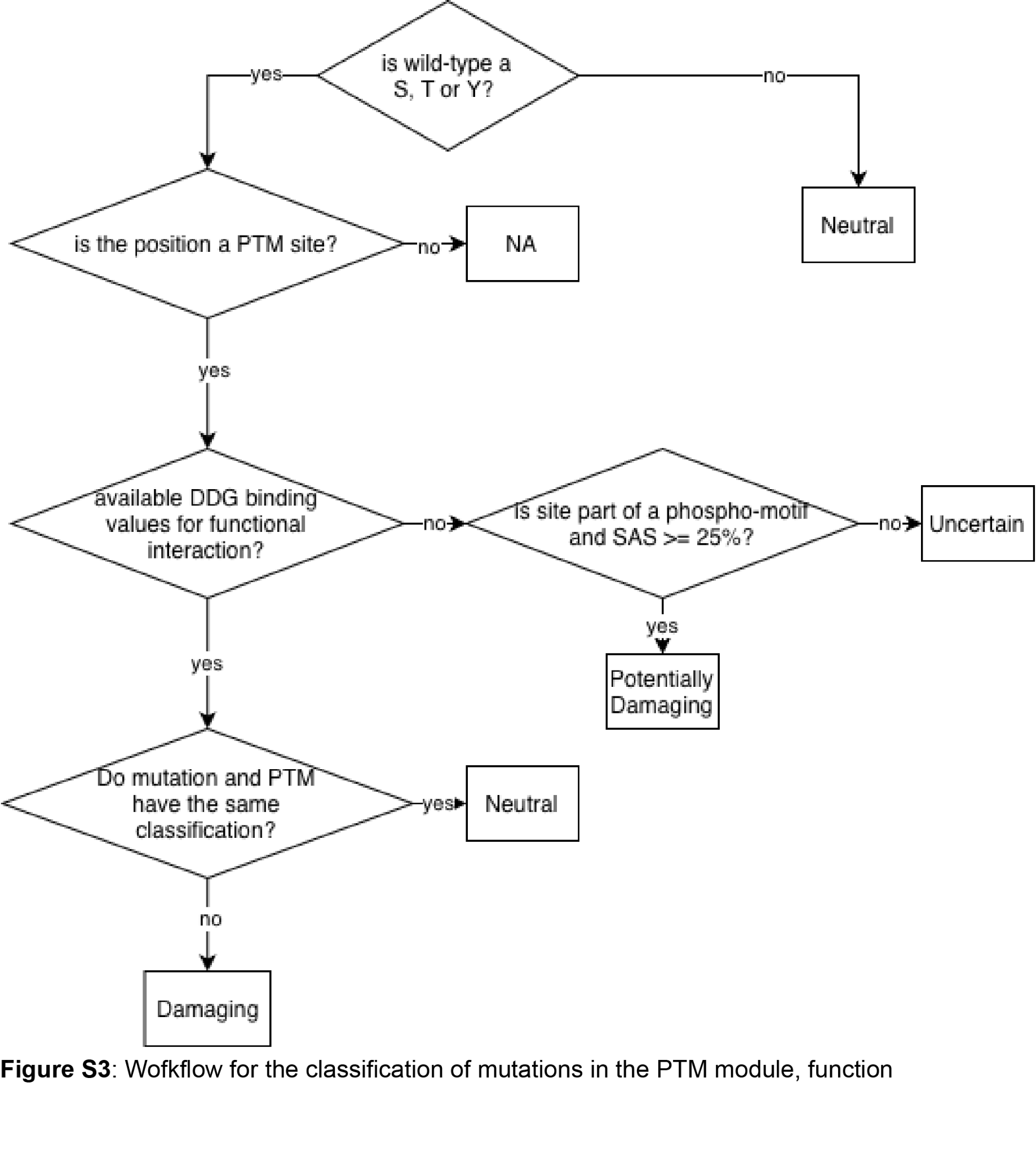

**Figure S3.3**. Workflow for the classification of mutations in the PTM module, function

*References*

[1] Tiberti, M., Di Leo, L., Vistesen, M. V., Kuhre, R. S., Cecconi, F., De Zio, D., & Papaleo, E. (2022). The Cancermuts software package for the prioritization of missense cancer variants: a case study of AMBRA1 in melanoma. *Cell Death & Disease*, *13*(10), 872.

[2] Hornbeck, P. V., Zhang, B., Murray, B., Kornhauser, J. M., Latham, V., & Skrzypek, E. (2015). PhosphoSitePlus, 2014: mutations, PTMs and recalibrations. Nucleic acids research, 43(D1), D512-D520.

[3] Hubbard,S.J.& Thornton, J.M. (1993), 'NACCESS', Computer Program, Department of Biochemistry and Molecular Biology, University College London

[4] Schymkowitz, J., Borg, J., Stricher, F., Nys, R., Rousseau, F., & Serrano, L. (2005). The FoldX web server: an online force field. Nucleic acids research, 33(suppl_2), W382-W388.

[5] Kumar, M., Michael, S., Alvarado-Valverde, J., Mészáros, B., Sámano‐Sánchez, H., Zeke, A., ... & Gibson, T. J. (2022). The eukaryotic linear motif resource: 2022 release. Nucleic acids research, 50(D1), D497-D508.
