## Supplementary Text S3 for "MAVISp: A Modular Structure-Based Framework for Protein Variant Effects"

**Supplementary Text S3. Benchmarking of thresholds for DeMaSk and GEMME scores against ClinVar interpretations.**

DeMaSk is a method for the prediction of the overall effect of mutations ([10.1093/bioinformatics/btaa1030](https://doi.org/10.1093%2Fbioinformatics%2Fbtaa1030)). It is based on a linear regression model that combines an asymmetric amino acid substitution matrix as well as evolutionary information. The most important outcome of DeMaSk is a delta fitness score (Δfitness), which can span negative to positive value, with negative values meaning the mutation is predicted to have loss-of-fitness effects, while positive for gain-of-fitness, and 0 for neutral. Nonetheless, the original authors of the method never specified a specific threshold to classify variants into neutral or non-neutral classes.

In this context, the following document provides an overview of the benchmarking process and results to evaluate the threshold to apply to DeMaSk scores within the MAVISp framework. In MAVISp, we classify a mutation according to Δfitness in two classes: Damaging, which includes both gain-of-fitness and loss-of-fitness cases, and Neutral. We therefore sought to find a single threshold for the DeMaSk score, to be applied in absolute value for both loss-of-fitness (negative score) and gain-of-fitness (positive score) intervals. In particular, we used loss-of-fitness information to identify a suitable threshold as the majority of our DeMaSk predictions had such classification (Δfitness < 0; see below), and because our original dataset is highly enriched of tumor suppressor proteins, for which a Damaging mutation would likely lead to loss-of-fitness. To do so, we developed a pipeline using R and Bash script (which can be found at https://www.github.com/ELELAB/MAVISp_pathogenicity_predictors_benchmark) for data mining, filtering and assessing predictive performances.

We used the 314 protein entries in the MAVISp database obtained on 13-02-2024 and we retained the variants that were annotated in ClinVar with a review status higher than or equal to 3. For each variant with an available ClinVar classification, we further processed it according to the internal MAVISp dictionary (available on the MAVISp GitHub, [clinvar_interpretation_internal_dictionary.txt](https://github.com/ELELAB/MAVISp/blob/main/mavisp/data/clinvar_interpretation_internal_dictionary.txt)) to obtain a final classification as Pathogenic or Benign. This was done to ensure handling dubious classifications in a predictable manner. This was the case, for instance, if any of the protein mutations was associated with multiple available ClinVar classifications. Finally, following this conversion step, we only retained variants that were classified as Pathogenic or Benign. We ended up with a total of 463 variants for analysis of which 373 pathogenic and 90 benign variants. The score distribution is reported in **Figure S4.1**. One should notice that in the data collected so far in the database, we observed more cases of negative scores (i.e., loss of fitness) and the distribution is skewed to the left. This could be due, as stated above, to the fact that a large majority of the proteins in the MAVISp database belong to the class of tumor suppressors and if they are mutated are expected to have a loss of function phenotype.

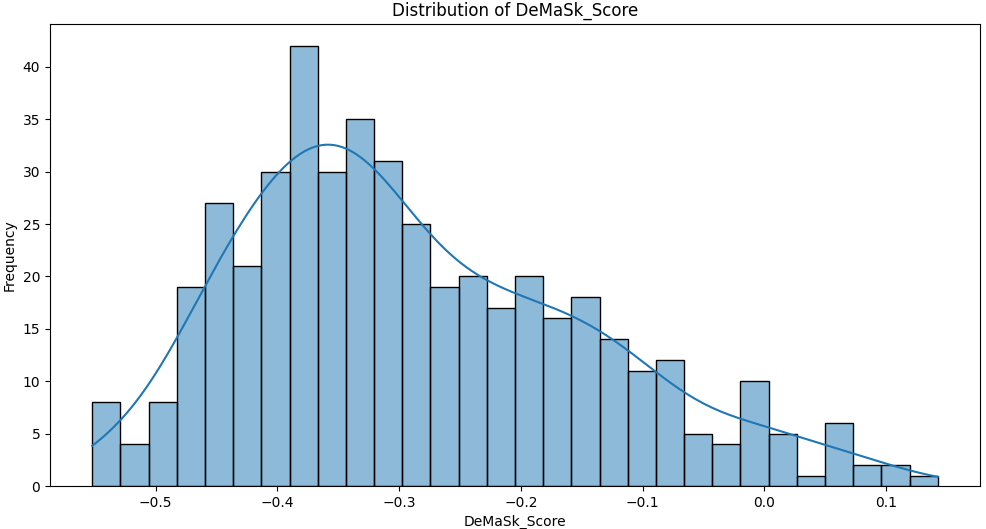

**Figure S4.1 Distribution of DeMaSk scores used in the analyses.** The figure consists of a bar plot,, illustrating the distribution of DeMaSk scores used in the study, and the density of data points calculated via kernel density estimation. In particular, the graph shows the frequency of DeMaSk scores predicted for mutations that had a ClinVar review status of 3 or 4.

As previously introduced, we aim to identify a threshold that would distinguish loss-of-fitness cases (here considered damaging) with respect to neutral cases. We generated a ROC curve to assess the performance of the prediction, illustrating the sensitivity-specificity trade-off across varying Δfitness thresholds. We obtained an AUC of 0.87 which falls within a range indicative of improved discrimination between positive and negative instances in the classification task (**Figure S4.2**).

Furthermore, we analyzed the classification performances with different thresholds, resulting in an optimal threshold of approximately -0.25 as the optimal trade-off between sensitivity and specificity. At this threshold, we obtained the smallest distance of 0.0685 between the ROC curve and point (1, 1) with a sensitivity of 0.77 and specificity of 0.86.

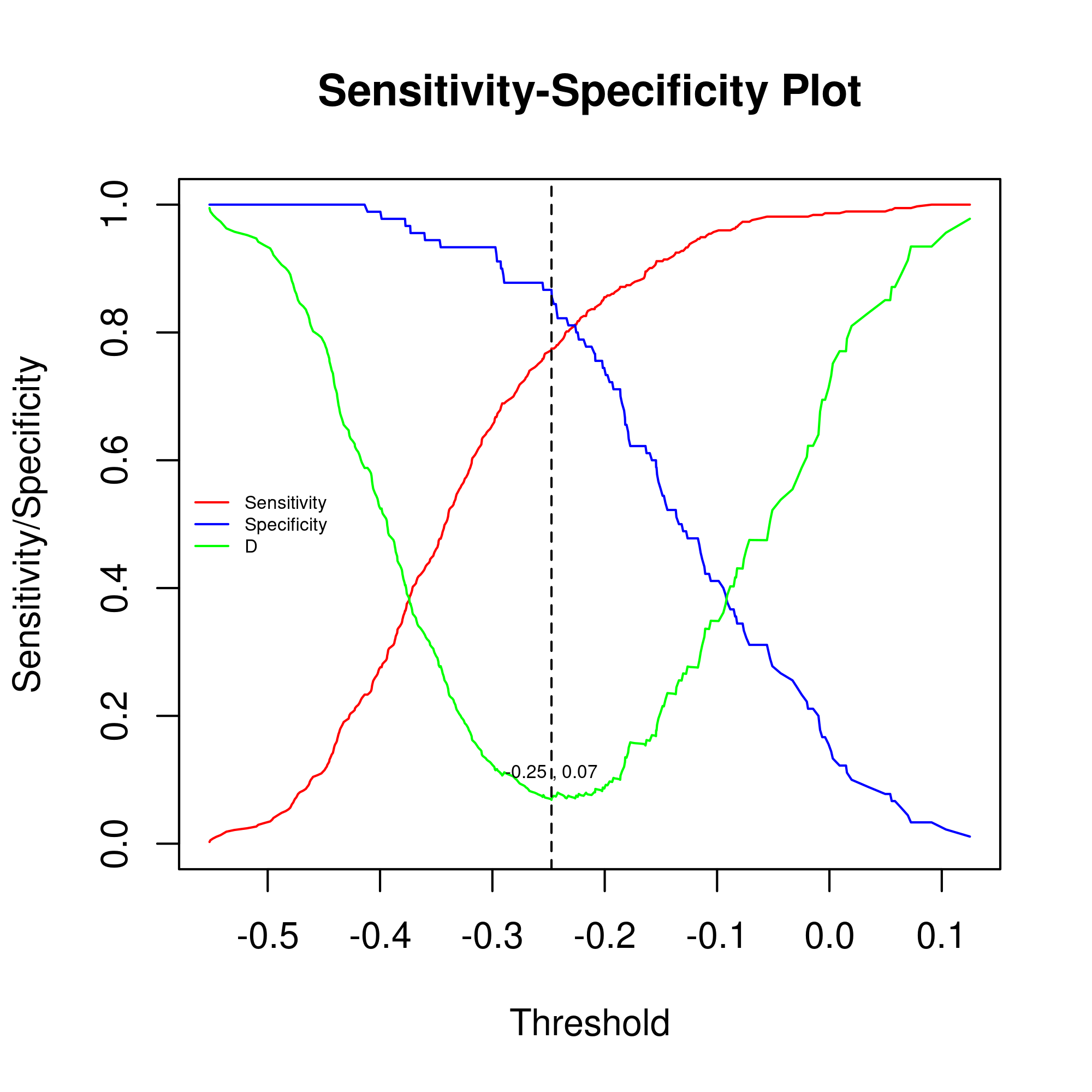

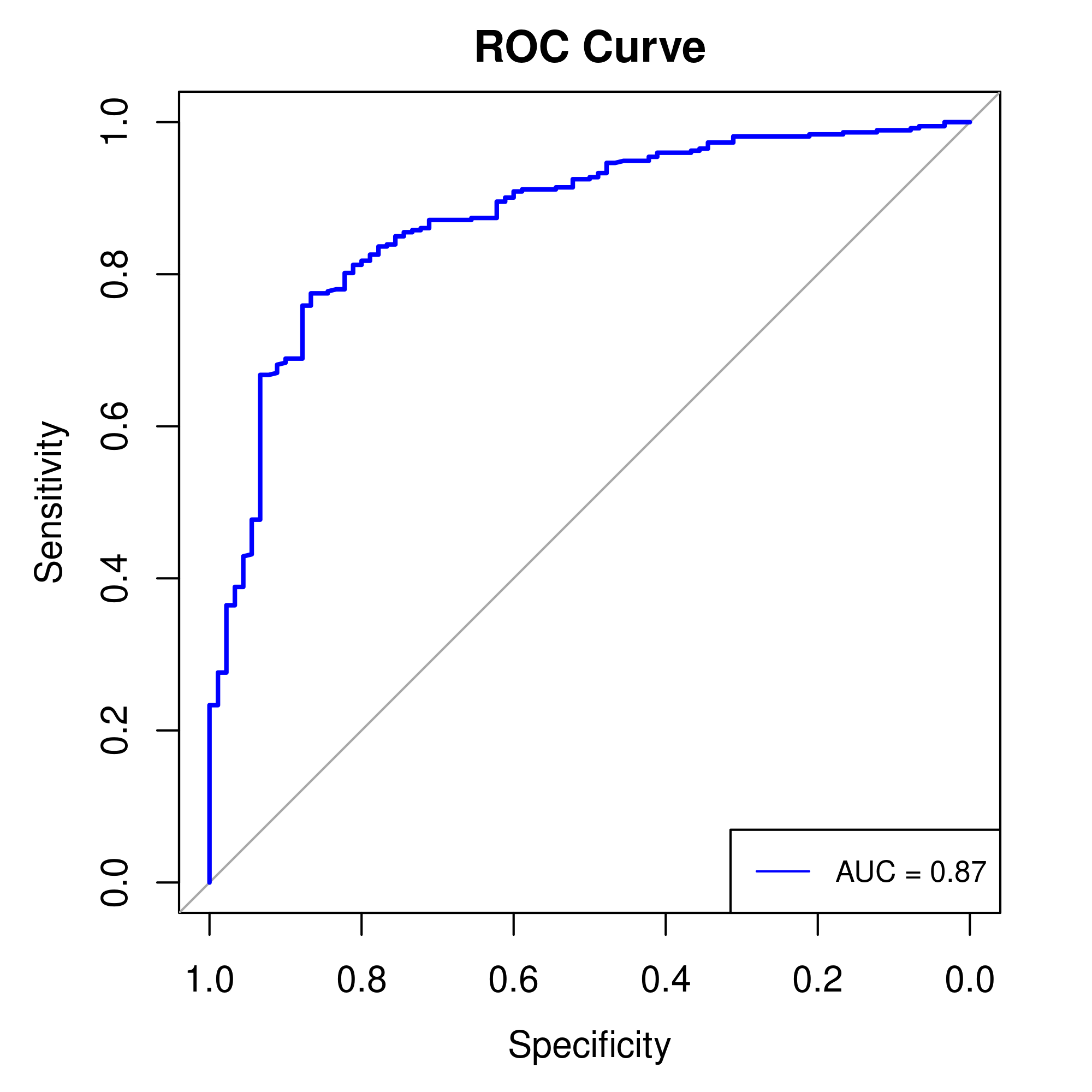

**Figure S4.2**: DeMaSk ROC and Sensitivity-Specificity Curves. A) The panel illustrates the ROC curve generated for DeMaSk using variants from the MAVISp database with ClinVar review status 3 or 4. B) The panel shows the sensitivity (red), specificity (blue) and the D function (green), representing classification performances for DeMaSk. D is the euclidean distance between the ROC curve and the (1, 1) point in the plot, which would represent optimal specificity and sensitivity. Lower D values indicate better performance. A vertical line marks the threshold with the lowest D.

We applied the same methodology to the analysis of GEMME (<https://doi.org/10.1093/molbev/msz179>) scores where negative values represent damaging mutations. In this case, we used 295 variants for which we had data in the MAVISp database and a clear classification for pathogenicity in ClinVar with review status of 3 or 4. Of these, 232 were pathogenic and 63 benign variants as reported in ClinVar according to the MAVISp internal dictionary. We obtained an AUC of 0.86 and an optimal threshold of -3. The results are reported in **Figure S4.3**.

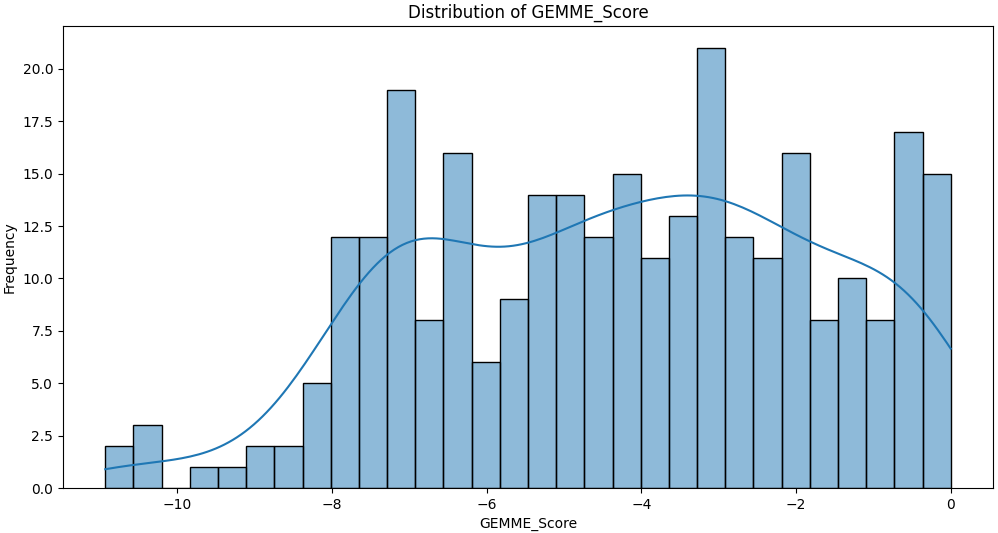

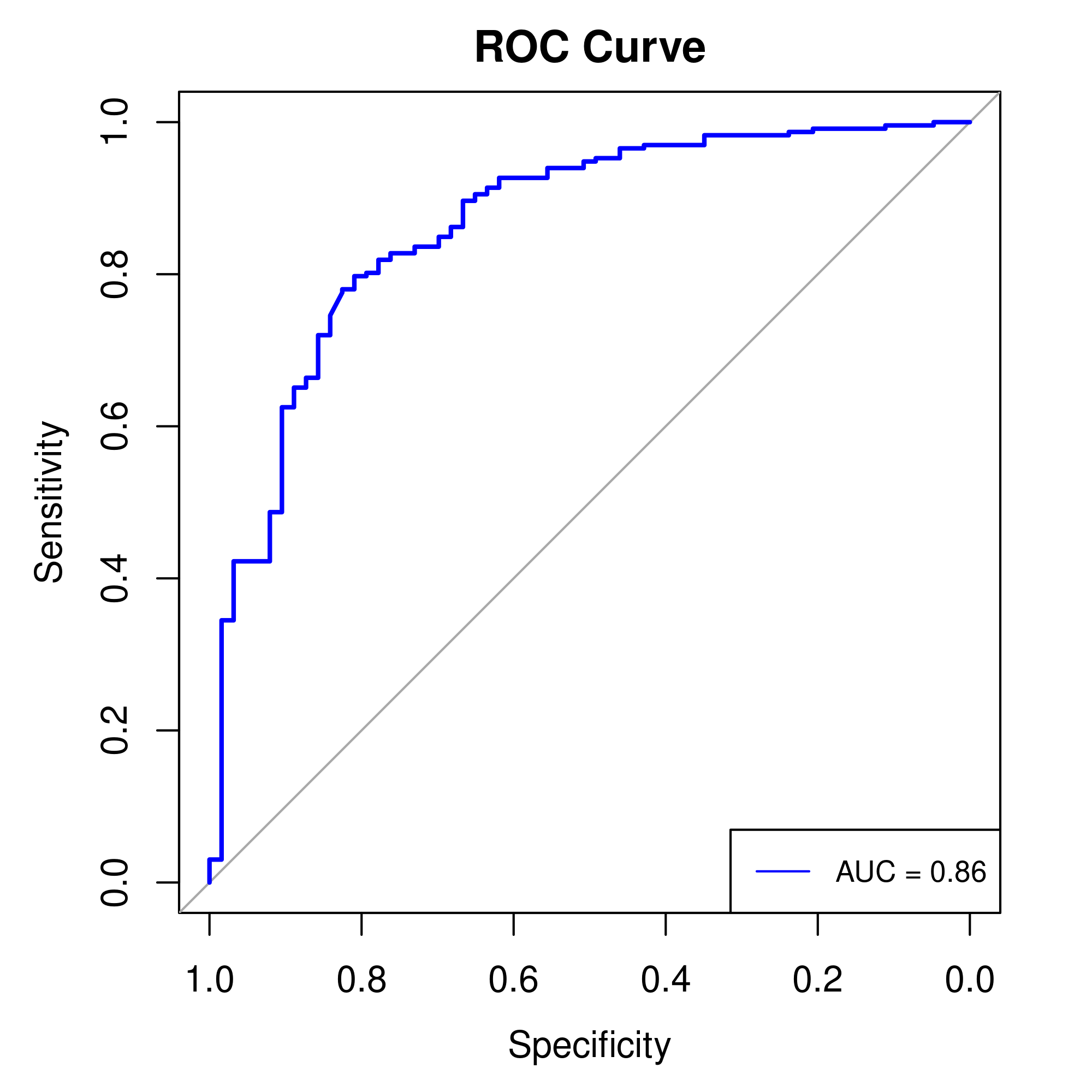

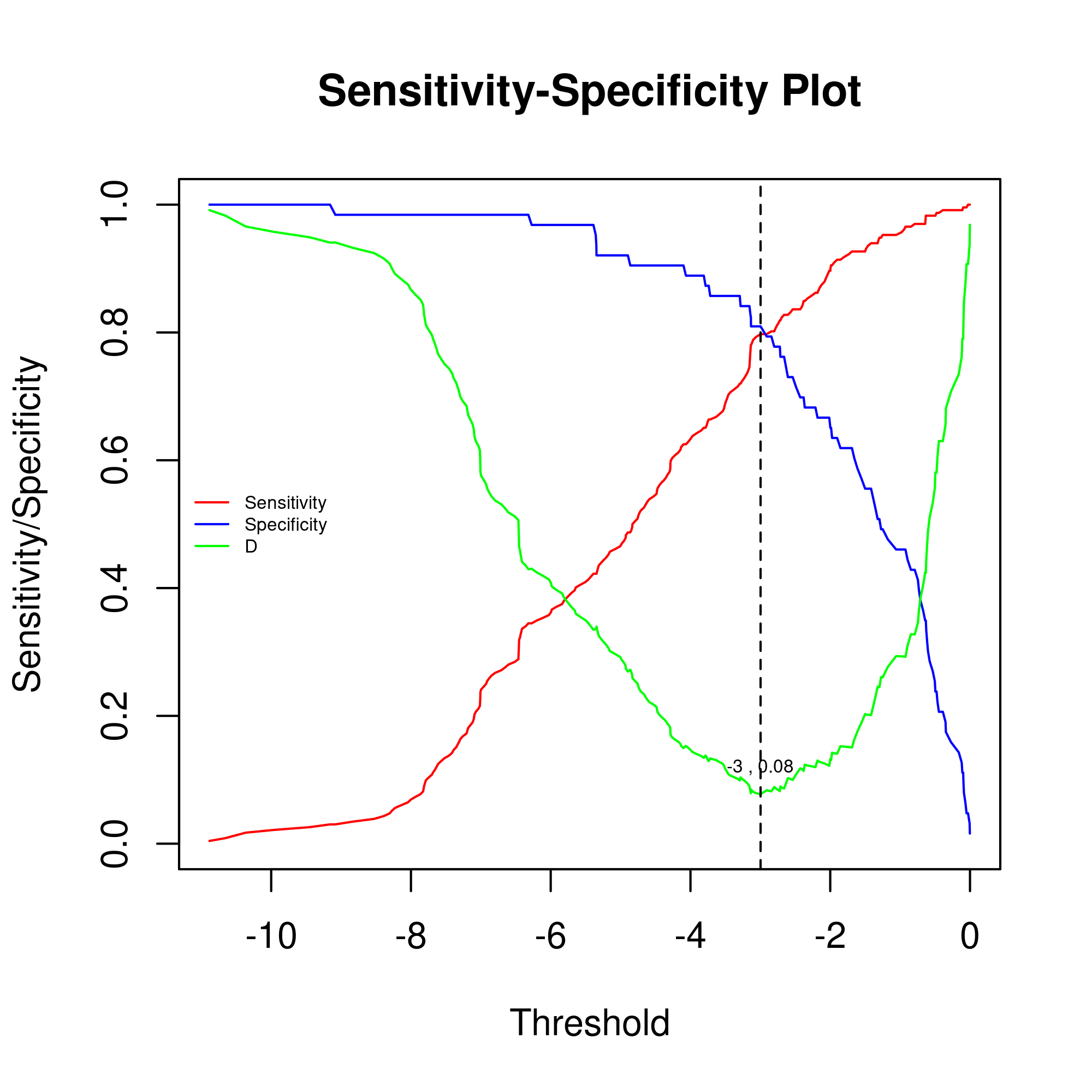

**Figure S4.3** Distribution plot, ROC and Sensitivity-Specificity Curves for GEMME. A) The panel illustrates the distribution of the GEMME scores, B) The panel shows the ROC curve generated for GEMME using variants from the MAVISp database with ClinVar review status 3 or 4. C) The panel shows the sensitivity (red), specificity (blue) and the D function (green), representing classification performances for GEMME. D is the euclidean distance between the ROC curve and the (1, 1) point in the plot, which would represent optimal specificity and sensitivity. Lower D values indicate better performance. A vertical line marks the threshold with the lowest D.
